## supplementary material for "Brain disconnections link structural connectivity with function and behaviour"

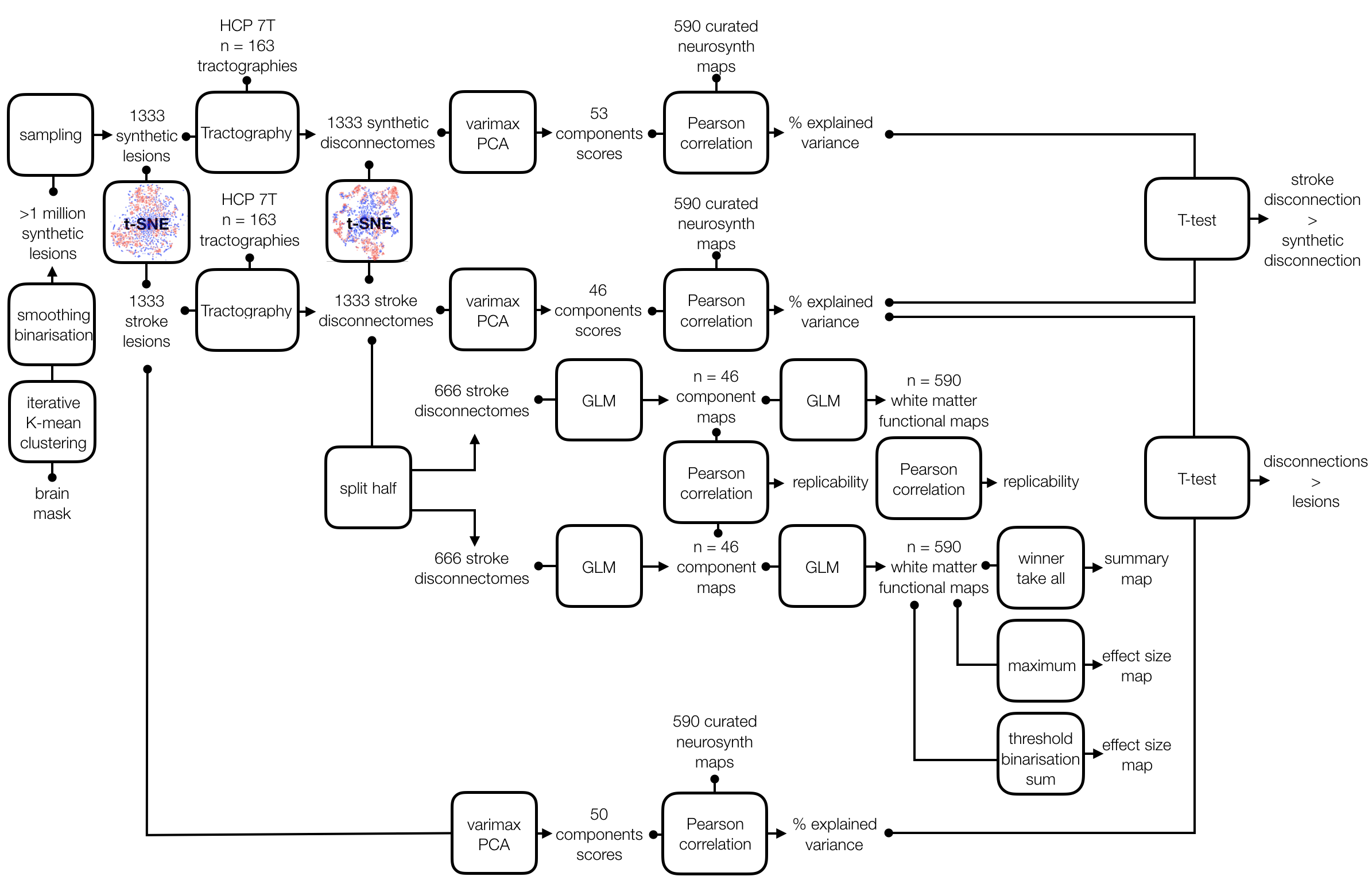

Supplementary figure 1: Graphical summary of the neuroimaging workflow

**A. Component maps**

Component maps correspond to the white matter structure subjacent to each component derived from a multiple regression between the component score (e.g. the independent variable) and the voxel probability of disconnection (e.g. the dependant variables) calculated in the two set of 666 stroke disconnectome maps. Replicability of these maps was assessed using a Pearson correlation between the results derived from the two set of 666 stroke disconnectome maps.

For each of the 46 components, we provide below an illustration including axial, coronal and sagittal display as well as a scatter plot of the highest correlation between the component score and task-related activation. The level of replicability of the component map is also reported together with an anatomical description discussed with regard to previously published lesion studies.

1) *1^st^ Component*

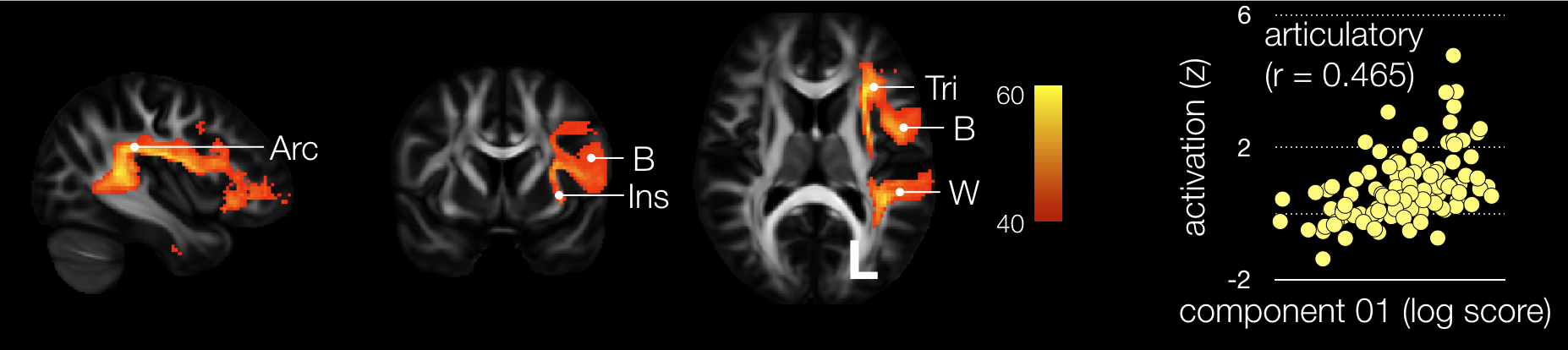

Supplementary figure 2: First component map side-by-side with its correlation with task related functional MRI (Pearson r > 0.202 are significant after Bonferroni correction for multiple comparisons). Arc: Arcuate fasciculus, B: Broca area, Ins: Insula, W: Wernicke area, Tri: pars triangularis

Component 1 correlated significantly with ‘*articulatory’* related task-*f*MRI activations (Pearson r = 0.465). Component 1 map was highly replicable (r = 0.975) and involved the arcuate fasciculus, which, together with a set of short U shape fibres in the frontal lobe, connected the Broca area and the pars triangularis in the frontal lobe, Wernicke area in the temporal lobe and the insula in the left hemisphere. Disconnection of the arcuate fasciculus as well as damage to the insula in stroke has been associated with impairment in speech production, particularly articulatory planning deficits ^1,2^. Importantly, a lesion of the superior temporal gyrus (named here Wernicke area) also manifest with some language production impairment ^3^ and is activated in healthy controls during the generation of noun and verbs ^4^. Hence behavioural manifestations occurring after a lesion to the connections and the areas significantly associated with component 1 confirm its involvement with ‘articulatory’ function.

2) *2^d^ Component*

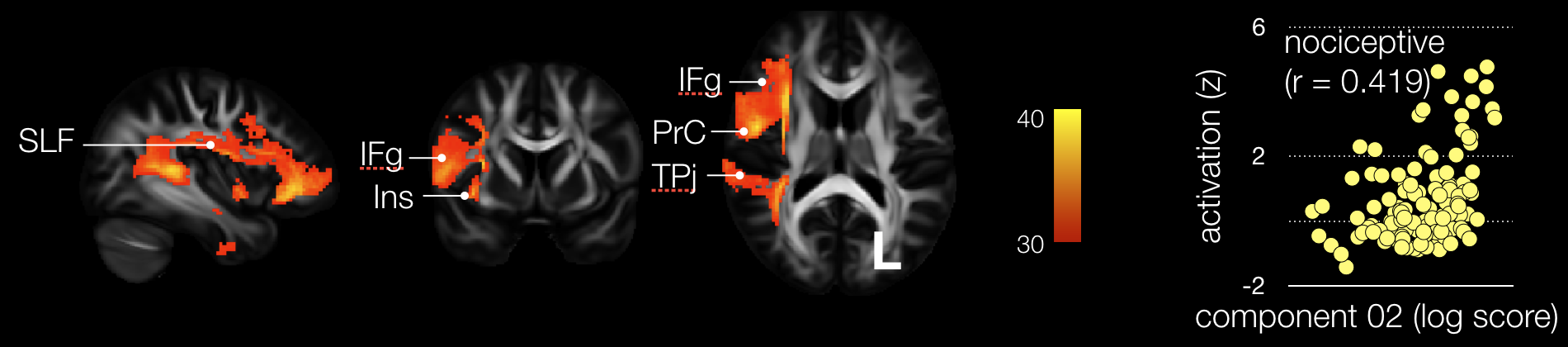

Supplementary figure 3: Second component map side-by-side with its correlation with task related functional MRI. SLF: Superior Longitudinal fasciculus, IFg: Inferior Frontal gyrus, Ins: Insula, TPj: Temporo-Parietal junction.

Component 2 correlated significantly with ‘*nociceptive’* related task-*f*MRI activations (r = 0.419). Component 2 map was highly replicable (r = 0.933) and involved the arcuate fasciculus, which, together with a set of short U shape fibres in the frontal lobe, connected the inferior frontal and precentral gyri, the temporo-parietal junction and the insula in the right hemisphere. Disconnection of the superior longitudinal fasciculus as well as damage to the insula and the precentral gyrus in stroke has been associated with disorder in body awareness, particularly anosognosia for hemiplegia that prevent patients to respond to stimuli on the controlesional side of their body and has an important role in associating internal emotional and bodily state signals with external sensory information^5^. Additionally, direct lesion of the insula lesion alters pain perception^6^. Thus direct lesion to the connection and the regions significantly associated with component 2 confirms its involvement in ‘nociception’.

3) *3^rd^ Component*

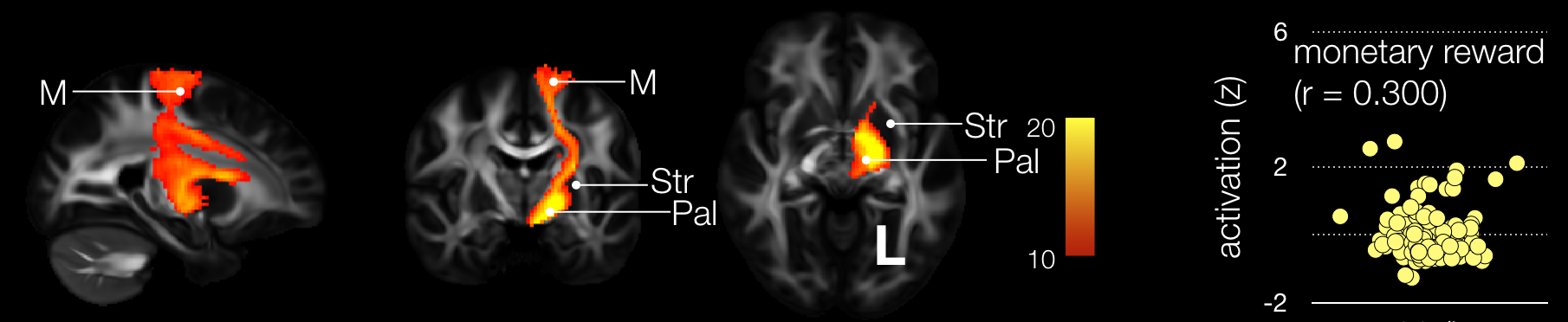

Supplementary figure 4: Third component map side-by-side with its correlation with task related functional MRI. M: Motor Cortex, Str: Striatum, Pal: Pallidum.

Component 3 correlated significantly with ‘*monetary reward’* related task-*f*MRI activations (r = 0.300). Component 3 map was well replicable (r = 0.857) and involved the striato-pallidal connections as well as projection of the motor cortex on the striatum in the left hemisphere. Direct lesion of the connections between the striatum and the pallidum has been shown to impact the relationship between reward and motor output^7^ and is thus compatible with the association between ‘monetary reward’ activation and component 3 map.

4) *4th Component*

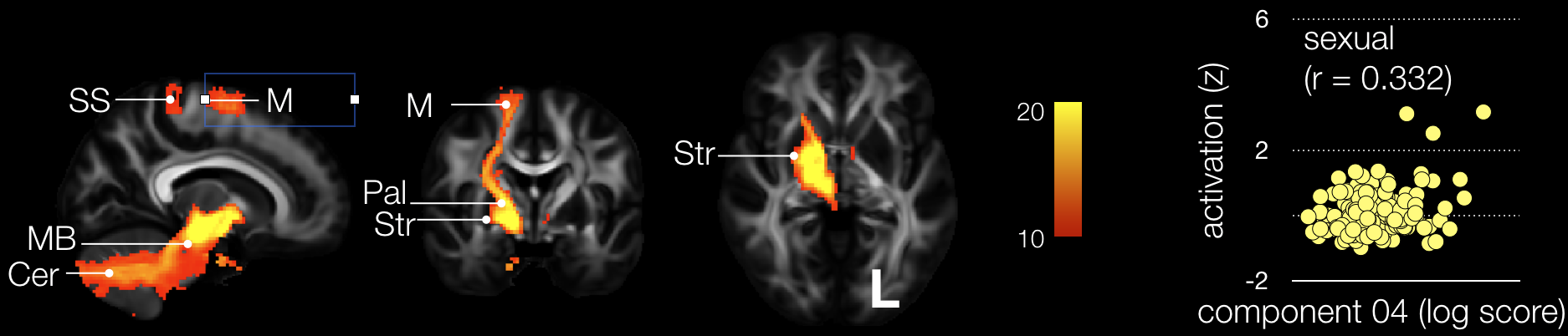

Supplementary figure 5: Fourth component map side-by-side with its correlation with task related functional MRI. M: Motor Cortex, SS: Somatosensory cortex, PreSMA: Pre-Supplementary area, Cer: Cerebellum, MB: Midbrain Str: Striatum, Pal: Pallidum.

Component 4 correlated significantly with ‘*sexual’* related task-*f*MRI activations (r = 0.332). Component 4 map was well replicable (r = 0.888) and mostly mirrored Component 3 map in the right hemisphere with a significant involvement of striato-pallidal connections as well as projection of the motor cortex on the striatum in the right hemisphere. Additionally Component 4 map involved the connections with the midbrain, the pre-supplementary area and the cerebellum. Lesions in the right hemisphere were associated with more difficulties in reaching orgasm than lesions in the left hemisphere ^8^ and patients with lesions in the striatum and the cerebellum report a decrease of sexual desire^9^. Non-human primates also activate their midbrain, particularly the periaqueductal gray when sexually aroused^10^ and genitals stimulation project on the medial portion of the somatosensory cortex in humans^11^. Hence, although sexuality in stroke has been poorly investigated, results derived from neurosurgery, neuroanatomy and high-field neuroimaging in non-human primates support the implication of component 4 map in ‘sexual’ related functions.

5) *5th Component*

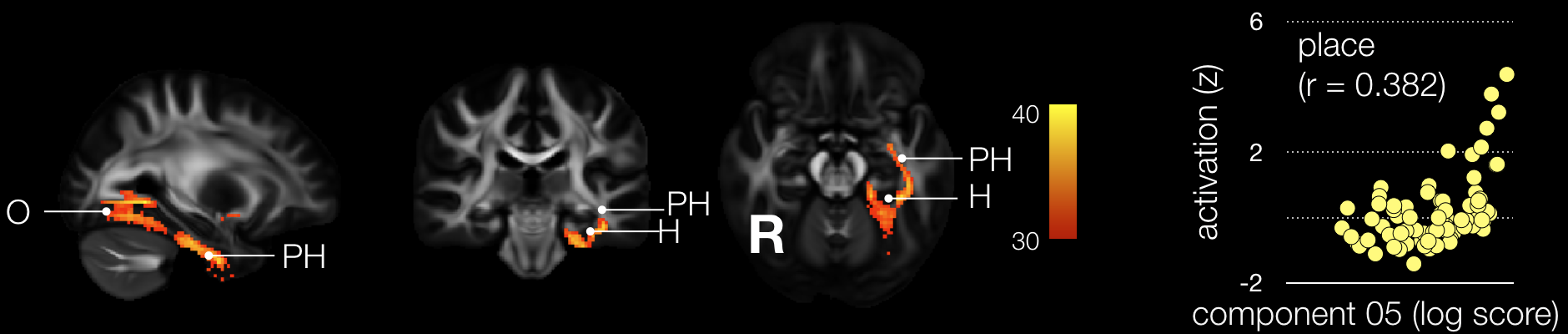

Supplementary figure 6: Fifth component map side-by-side with its correlation with task related functional MRI. O: Occipital Cortex, PH: Parahippocampus, H: hippocampus

Component 5 correlated significantly with ‘*place’* related task-*f*MRI activations (r = 0.382). Component 5 map was highly replicable (r = 0.903) and significantly disconnected the medial portion of the occipital lobe together with the inferior para-hippocampal gyrus and the hippocampus in the left hemisphere. While disorientation is a frequent symptom in stroke ^12^ the lesion location leading to it remains elusive. Notwithstanding this gap in the literature, a consensus emerge that the neural components of topographical representation should involve ventral occipitotemporal cortex and hippocampus (For a review see ^13^). All in all, previous results derived from neuroimaging in human and direct recordings in animal studies are compatible with Component 5 being associated to ‘place’-related functional MRI tasks.

6) *6th Component*

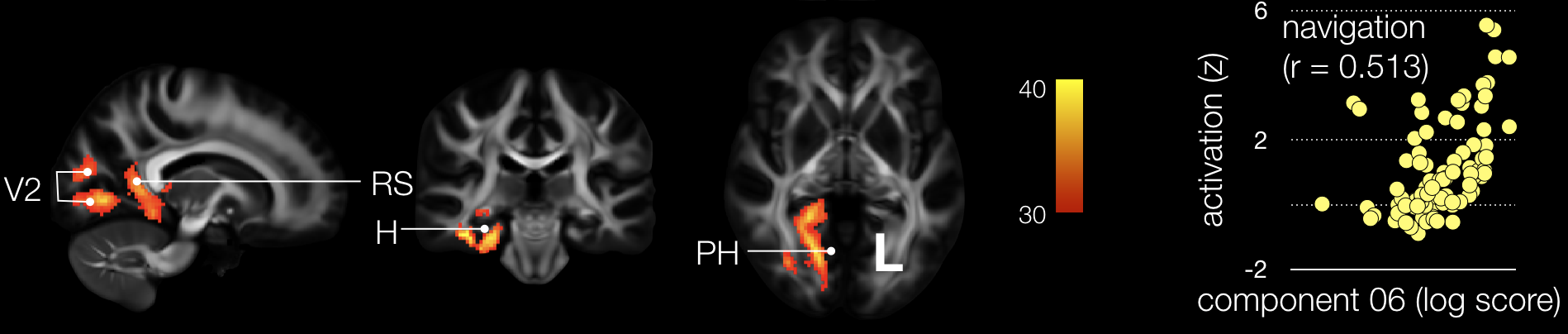

Supplementary figure 7: Sixth component map side-by-side with its correlation with task related functional MRI. V2: Secondary visual area, RS: Retrosplenial cortex, H: hippocampus, PH: Parahippocampal gyrus

Component 6 correlated significantly with ‘*navigation’* related task-*f*MRI activations (r = 0.513). Component 6 map was highly replicable (r = 0.927) and significantly disconnected the secondary visual areas the parahippocampal gyrus and the hippocampus as well as the retroplenial cortex in the right hemisphere. Stroke patients often show navigation impairment ^14^. The brain navigation system is well known to involve the posterior portion of the cingulum that connects the right hippocampus ^15^ to the retrosplenial cortex ^16,17^. Additionally, the ventral stream linking visual areas to the hippocampus parahippocampus and perirhinal areas is implicated for visual memory ^18^ critical to the success in any task of navigation. Hence the brain areas associated with component 6 are compatible with its involvement in ‘navigation’ brain processes.

7) *7th Component*

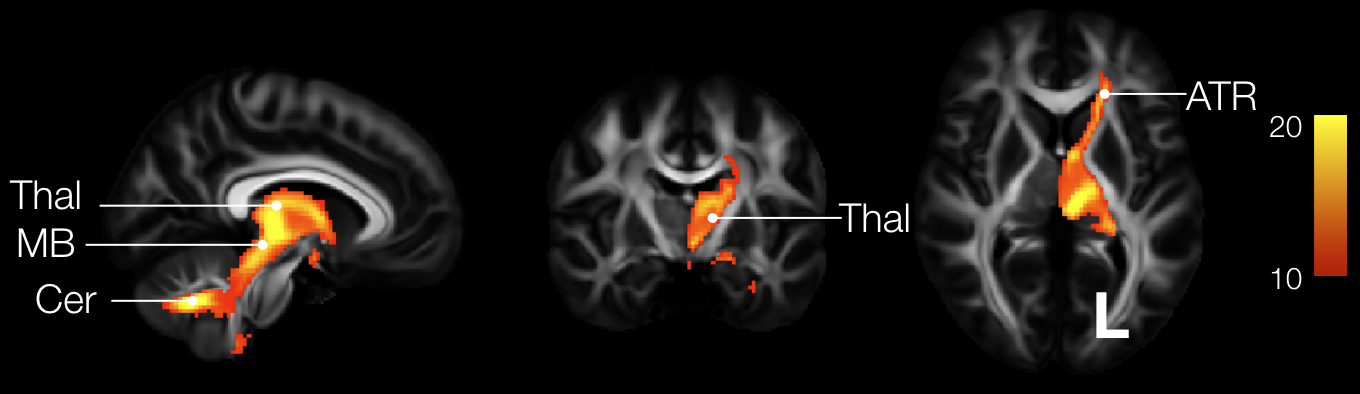

Supplementary figure 8: Seventh component map. Cer: Cerebellum, MB: Midbrain, Thal: Thalamus, ATR: Anterior Thalamic Radiations.

Component 7 map was highly replicable (r = 0.905) and significantly disconnected the cerebellum, midbrain, thalamus and anterior thalamic radiations in the left hemisphere. Correlations between component 7 and task-related functional activation did not survive Bonferroni correction for multiple comparisons (n = 590 functions).

8) *8th Component*

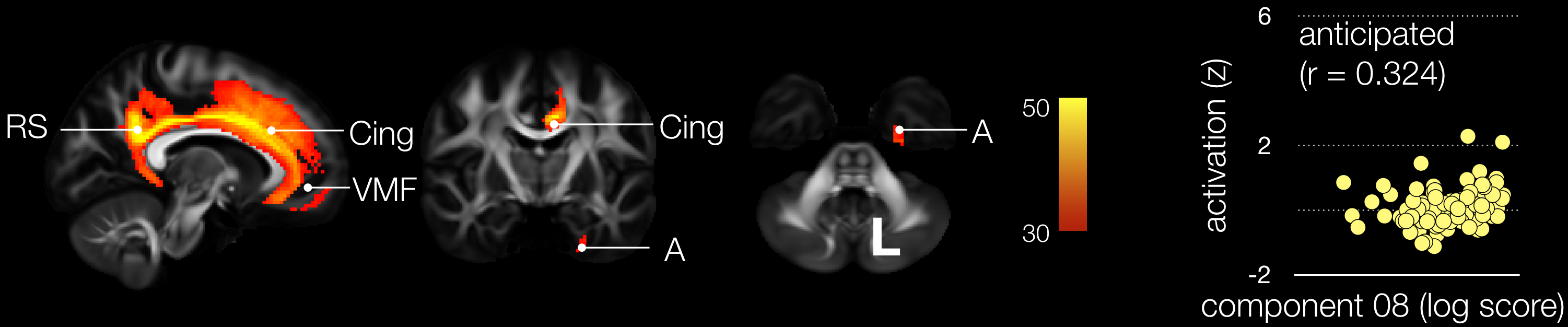

Supplementary figure 9: Eighth component map side-by-side with its correlation with task related functional MRI. RS: Retrosplenial cortex, Cing: Cingulum, VMF: Ventromedial frontal cortex, A: Amygdala

Component 8 correlated significantly with ‘anticipated’ related task-*f*MRI activations (r = 0.324). Component 8 map was highly replicable (r = 0.952) and significantly disconnected the ventromedial frontal cortex, the cingulum, the retrosplenial cortex and the amygdala in the left hemisphere. Direct lesions as well as disconnection of the ventromedial frontal cortex and amygdala in rats have been associated with an altered learning and representation of outcomes ^19,20^ and activations in the retrosplenial cortex in humans have been reported during tasks related to the planning of the future ^21^. Importantly, the cingulum white matter tract connects the ventromedial prefrontal cortex with the retrosplenial cortex and the amygdala ^22^. Therefore, results derived from lesion in rats and task activation in humans support the involvement of component 8 in ‘anticipatory’ processes.

9) *9th Component*

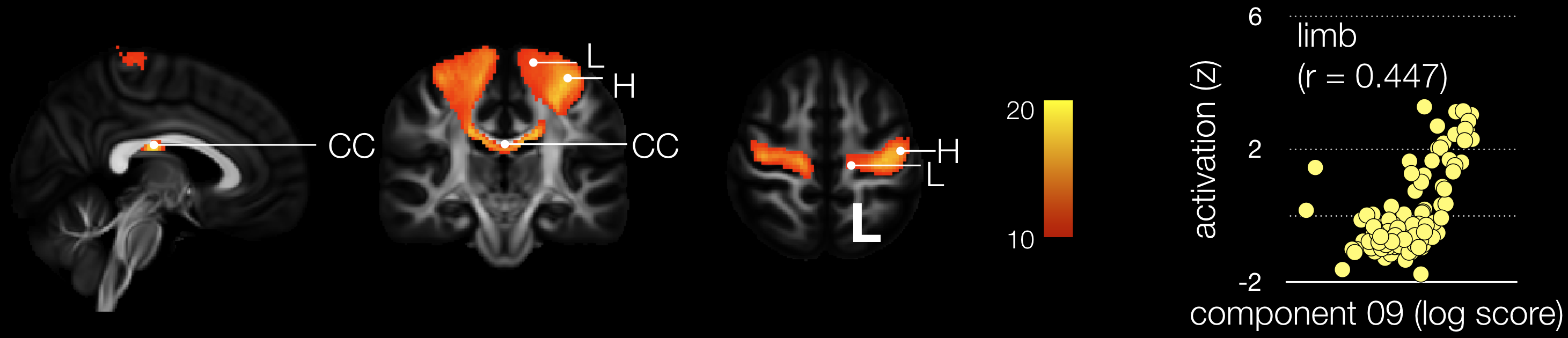

Supplementary figure 10: Ninth component map side-by-side with its correlation with task related functional MRI. CC: Corpus Callosum, L: Motor leg area, H: Motor hand area.

Component 9 correlated significantly with ‘limb’ related task-*f*MRI activations (r = 0.447). Component 9 map was moderately replicable (r = 0.782) and significantly disconnected the motor hand and leg areas bilaterally as well as the motor portion of the corpus callosum. Direct damage to the ‘middle knee’ of the central sulcus is well known to cause motor weakness in the contralateral hand ^23^, while lesion or direct per-operative electrical stimulation of the medial portion central sulcus are more likely to produce a disturbance of the lower limb ^24^. Importantly coordination disorders and intermanual conflict (i.e. alien hand syndrome or diagonistic apraxia) ^25,26^ and involuntary goal-directed movements of one hand (i.e. anarchic hand syndrome) ^27,28^ have been reported after a disconnection of the interhemispheric communication between the left and the right motor cortices. Hence lesions studies reported in the literature are compatible with the involvement of component 9 in ‘limb’ related processes and particularly their coordination.

10) *10th Component*

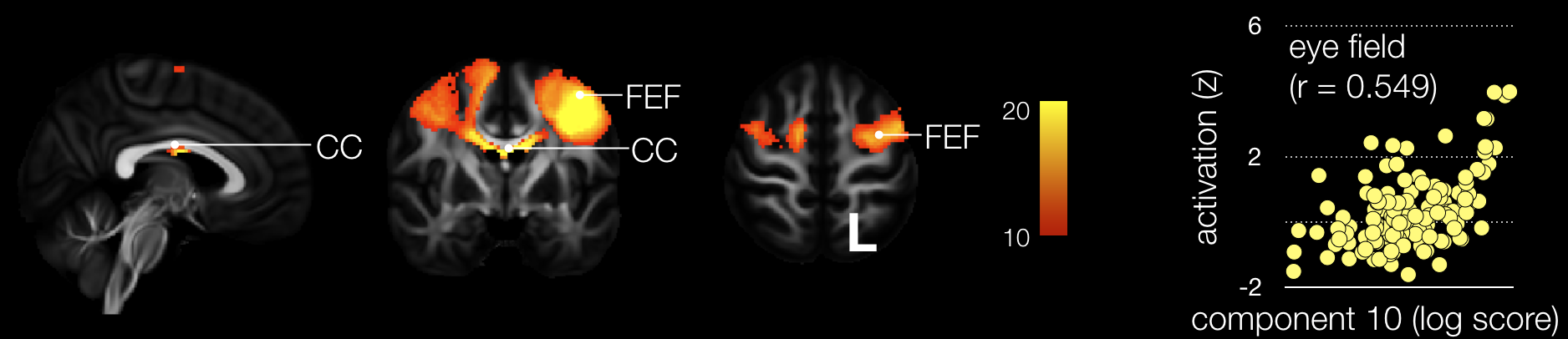

Supplementary figure 11: Tenth component map side-by-side with its correlation with task related functional MRI. CC: Corpus Callosum, FEF: Frontal Eye Field

Component 10 correlated significantly with ‘eye field’ related task-*f*MRI activations (r = 0.549). Component 10 map was moderately replicable (r = 0.786) and significantly involved the frontal eye field bilaterally as well as a portion of the corpus callosum. Irrepressible deviation of the gaze of both eyes have been reported after a stroke ^29^ or unilateral electrical stimulation ^30^ of the frontal eye field and is compatible with the implication of component 10 in ‘eye field’ function.

11) *11th Component*

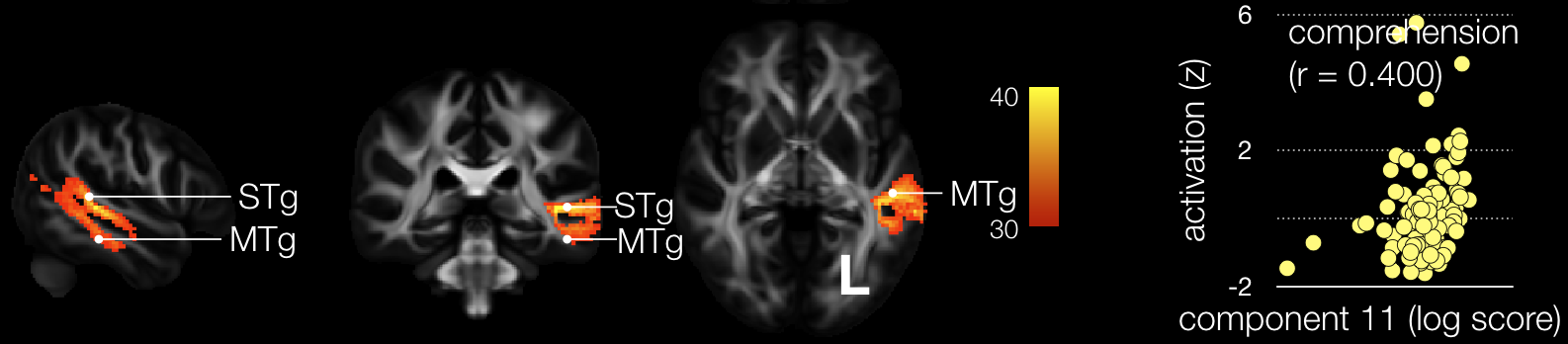

Supplementary figure 12: Eleventh component map side-by-side with its correlation with task related functional MRI. STg: Superior Temporal gyrus, MTg: Middle Temporal gyrus.

Component 11 correlated significantly with ‘comprehension’ related task-*f*MRI activations (r = 0.400). Component 11 map was highly replicable (r = 0.925) and significantly involved the disconnection of the superior temporal gyrus and the posterior part of the middle temporal gyrus in the left hemisphere. a lesion to the posterior part of the middle temporal gyrus or the superior temporal gyrus are associated with oral comprehension deficits ^31^. Further the importance of the connections in the depth of the temporal love have been recently highlighted for comprehension deficits (i.e. Wernicke aphasia) in stroke ^31^ and neurodegenerative disorders ^32^ confirming the hypothesis that Wernicke aphasia might well be a disconnection syndrome ^33^. Thus, the implication of component 11 in ‘comprehension’ processes is concordant with previous report in stroke and neurodegenerative diseases.

12) *12th Component*

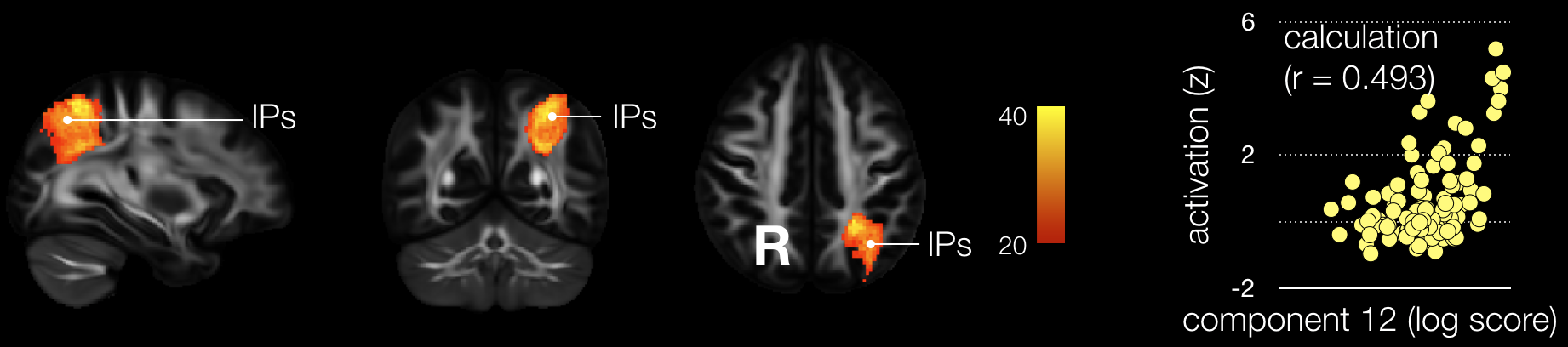

Supplementary figure 13: Twelfth component map side-by-side with its correlation with task related functional MRI. IPs: Intraparietal sulcus

Component 12 correlated significantly with ‘calculation’ related task-*f*MRI activations (r = 0.493). Component 12 map was highly replicable (r = 0.917) and significantly disconnected the intraparietal sulcus and its subjacent short white matter pathways in the left hemisphere. Per operative electrical stimulation ^34^ or direct lesion ^35^ of the white matter in the depth of the intraparietal sulcus of the left hemisphere has been reported as leading to the Gerstmann syndrome characterized by a quartet of symptoms including acalculia, agraphia, left–right disorientation, and finger agnosia ^36,37^. Therefore the involvement of component 12 in ‘calculation’ processes is compatible with previous neurosurgical and neurological reports.

13) *13th Component*

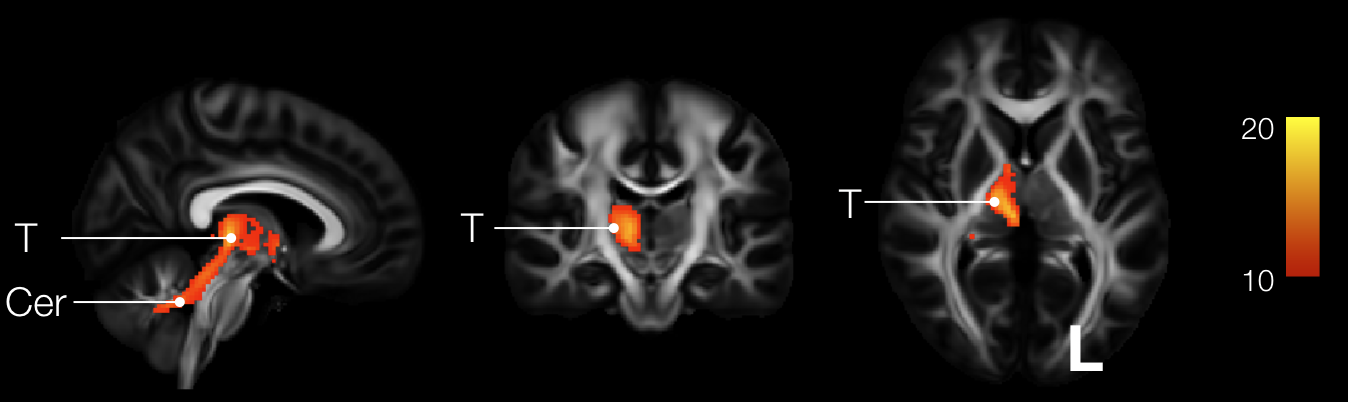

Supplementary figure 14: Thirteenth component map. T: Thalamus, Cer: Cerebellum.

Component 13 map was poorly replicable (r = 0.690) and significantly disconnected the cerebellum and the thalamus in the right hemisphere. Correlations between Component 13 and task-related functional activation did not survive Bonferroni correction for multiple comparisons (n = 590 functions).

14) *14th Component*

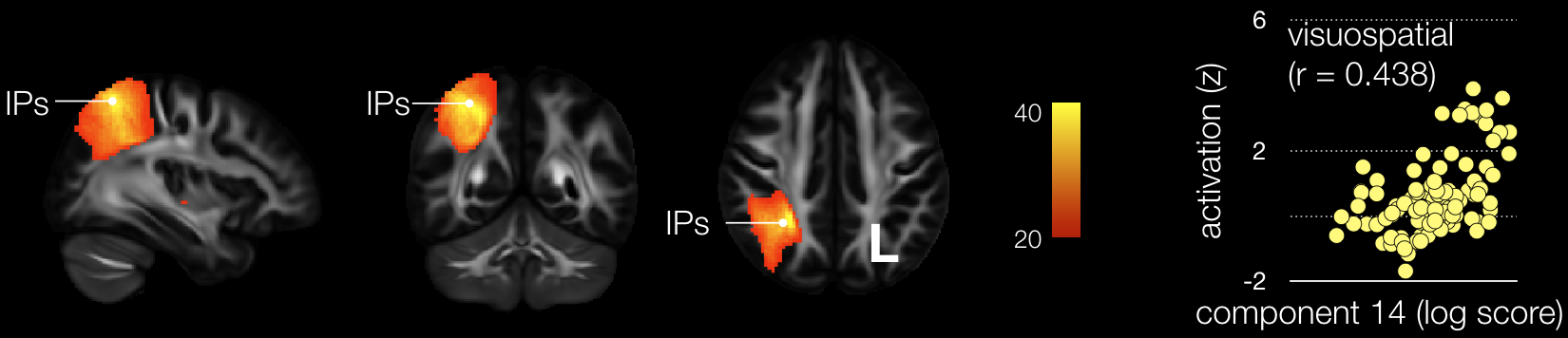

Supplementary figure 15: Fourteenth component map side-by-side with its correlation with task related functional MRI. IPs: Intraparietal sulcus.

Component 14 correlated significantly with ‘visuospatial’ related task-*f*MRI activations (r = 0.438). Component 14 map was highly replicable (r = 0.913), mirrored component 12 in the right hemisphere and significantly disconnected the intraparietal sulcus. Direct electrical stimulation ^38^ as well as lesions in the vicinity or in the white matter subjacent to the intraparietal sulcus ^39^ leads to signs of visuospatial neglect in humans and monkeys ^40^. Hence, results derived from lesion studies and per operative direct electrical stimulation in humans and monkeys are compatible with the implication of component 14 in ‘visuospatial’ processes.

15) *15th Component*

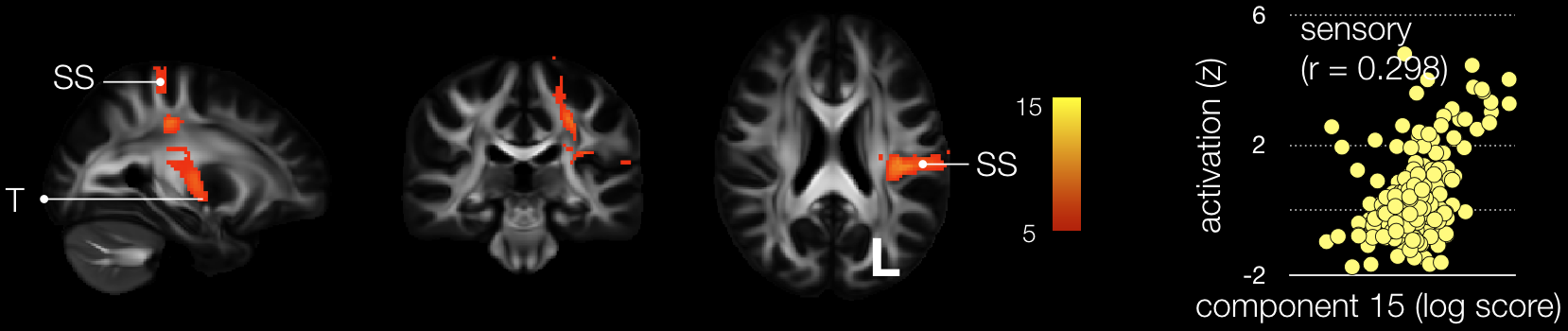

Supplementary figure 16: Fifteenth component map side-by-side with its correlation with task related functional MRI. SS: Somatosensory cortex, T: Thalamus

Component 15 correlated significantly with ‘sensory’ related task-*f*MRI activations (r = 0.298). Component 15 map was well replicable (r = 0.891) and significantly disconnected primary sensory areas from thalamus input in the left hemiphere. Disconnection of the thalamic radiations connecting the thalamus to the somatosensory cortex have been reported as leading to proprioceptive somatosensory symptoms ^41^. Thus, lesion mapping in patients showing somatosensory deficits confirm the involvement of component 15 in ‘sensory’ processes.

16) *16th Component*

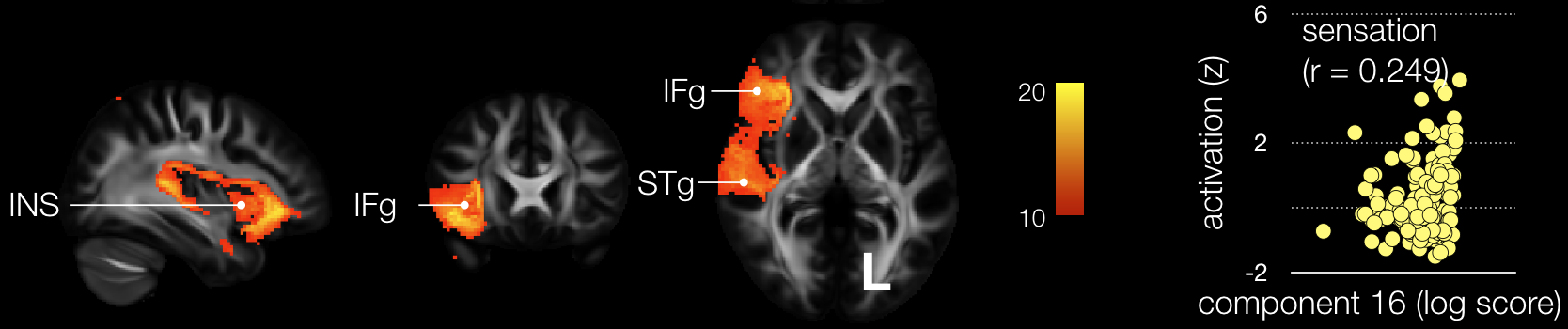

Supplementary figure 17: Sixteenth component map side-by-side with its correlation with task related functional MRI. SLF: Superior Longitudinal Fasciculus, INS: Insula, IFg: Inferior Frontal gyrus, STg: Superior Temporal gyrus

Component 16 correlated significantly with ‘sensation’ related task-*f*MRI activations (r = 0.298). Component 16 map was moderately replicable (r = 0.798) and disconnected significantly the inferior frontal gyrus, insula and superior temporal gyrus and their subjacent short white matter interconnections in the right hemisphere. While a visceral sensation and emotions role of the connections short white matter insular connections has been suggested ^42^, direct lesion to the insula in the right hemisphere affects patients’ ability to associate motor predictions with sensorimotor feedback ^43^. Hence previous anatomical studies and lesion studies seems compatible with the involvement of component 16 in ‘sensation’ brain processes.

17) *17th Component*

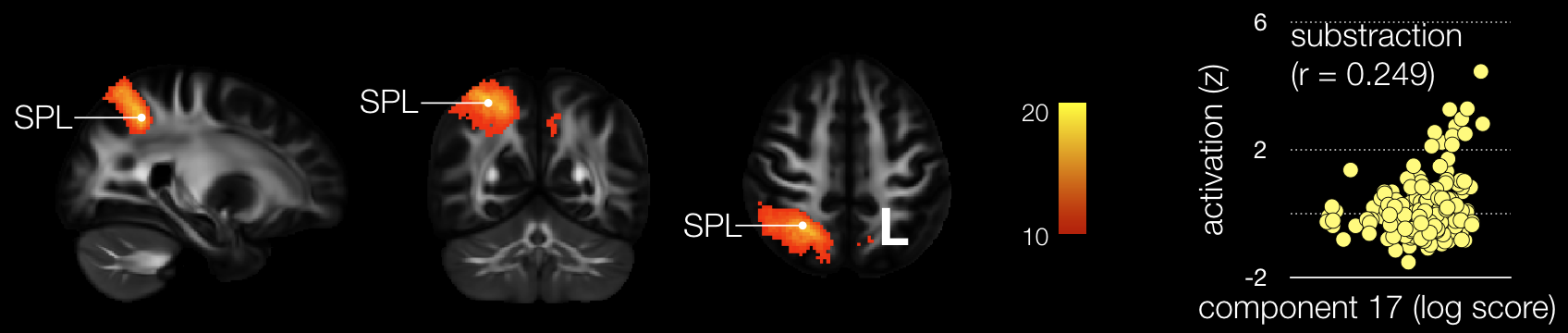

Supplementary figure 18: Seventeenth component map side-by-side with its correlation with task related functional MRI. SPL: Superior Parietal Lobule

Component 17 correlated significantly with ‘substraction’ related task-*f*MRI activations (r = 0.298). Component 17 map was moderately replicable (r = 0.725) and significantly disconnected significantly the right superior parietal lobule. While acalculia is usually located in the left hemisphere, several reports of acalculia after a lesion in the right parietal lobe exist ^44-46^ and concerned specifically writing arabic numerals to dictation, mental one-digit multiplication and mental one-digit substraction ^47^. Therefore the observation of stroke patients with a lesion in the right hemisphere is compatible with the involvement of component 17 in ‘substraction’ brain processes.

18) *18th Component*

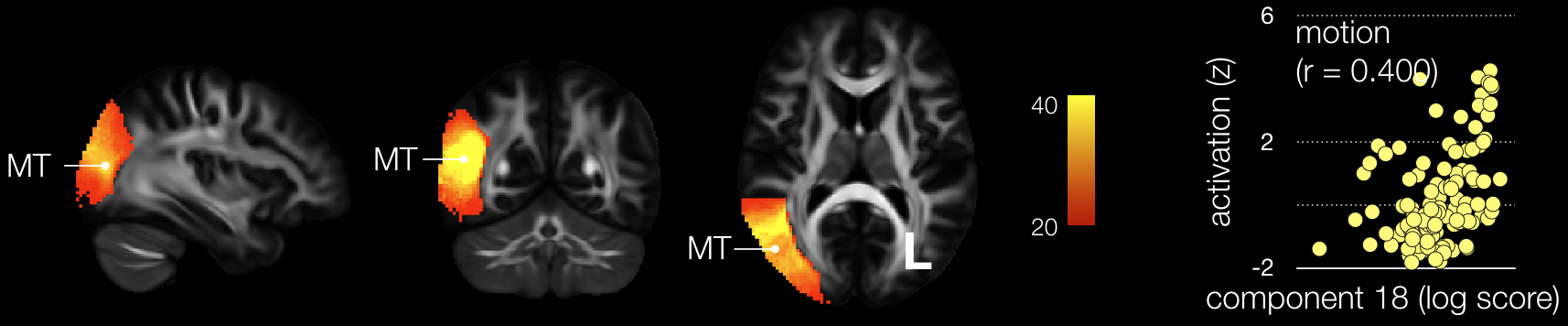

Supplementary figure 19: eighteenth component map side-by-side with its correlation with task related functional MRI. MT: middle temporal

Component 18 correlated significantly with ‘motion’ related task-*f*MRI activations (r = 0.298). Component 18 map was well replicable (r = 0.883) and significantly disconnected the MT area and it’s subjacent white matter in the right hemisphere. Deficits at the motion coherence task have been reported in cases of direct lesion or disconnection of MT ^48,49^. Hence, neuropsychological investigation in stroke patients appears to corroborate the implication of component 18 in ‘motion’ brain processes.

19) *19th Component*

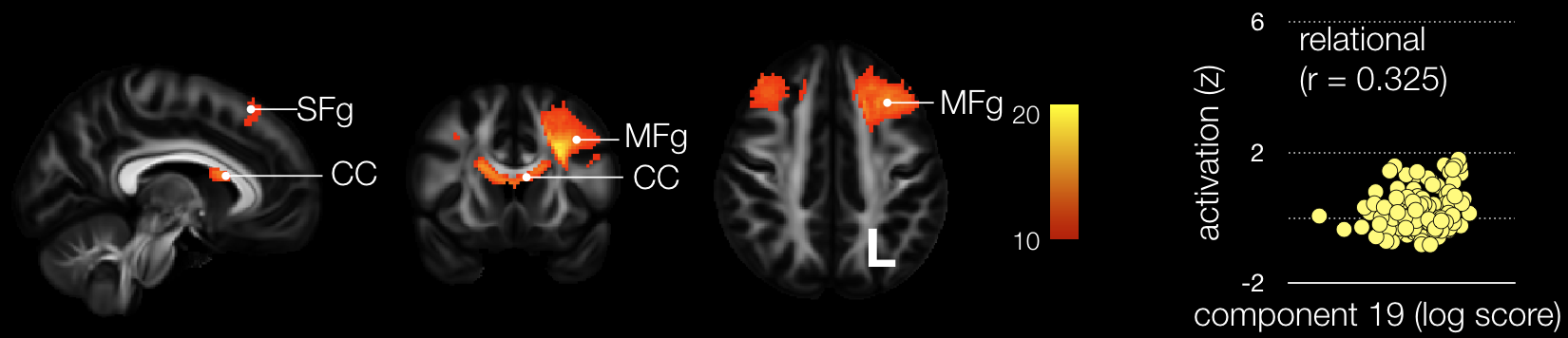

Supplementary figure 20: Nineteenth component map side-by-side with its correlation with task related functional MRI. SFg: Superior Frontal gyrus, MFg: Middle Frontal gyrus, CC: Corpus Callosum.

Component 19 correlated significantly with ‘relational’ related task-*f*MRI activations (r = 0.298). Component 19 map was well replicable (r = 0.825) and significantly disconnected the middle frontal gyrus bilaterally as well as its callosal connection. In the left hemisphere, a small involvement of the presupplementary area was also visible. Relational deficits have been previously reported after a stroke in the frontal lobe ^50^ particularly in the anterior portion of the middle frontal gyrus and its subjacent connections ^51^. However, there is no report to our knowledge of lesion or disconnection of the superior frontal gyrus associated with relational deficits. Hence while the lesion and disconnection of the middle frontal gyrus support the involvement of component 19 in ‘relational’ brain processes, the involvement of the superior frontal gyrus in these processes remains to be explored.

20) *20th Component*

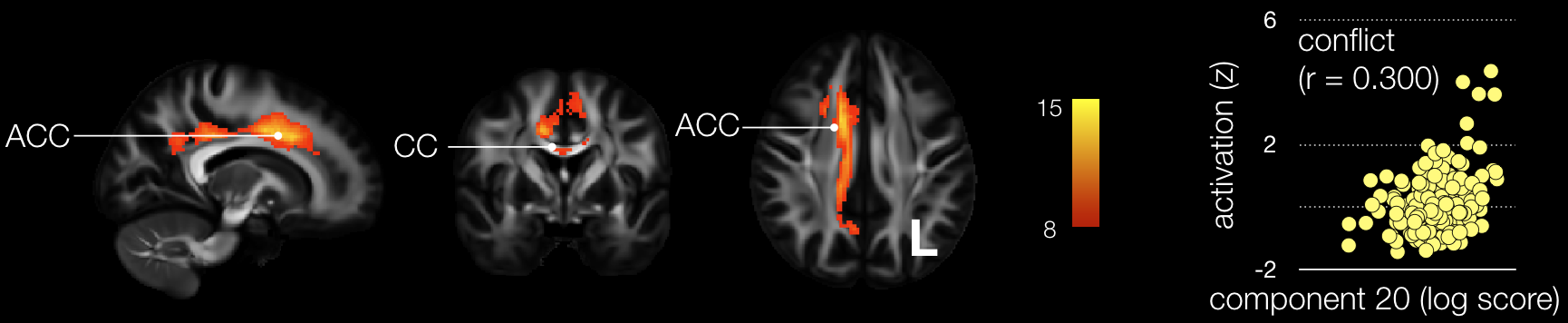

Supplementary figure 21: Twentieth component map side-by-side with its correlation with task related functional MRI. ACC: Anterior cingulate cortex, CC: Corpus Callosum.

Component 20 correlated significantly with ‘conflict’ related task-*f*MRI activations (r = 0.300). Component 20 map was well replicable (r = 0.877) and significantly disconnected the right anterior cingulate cortex and its connections with the corpus callosum. Lesions in the anterior cingulate cortex has been significantly associated with the absence of conflict effect (i.e. Simon effect) ^52^. Hence, stroke studies confirm the involvement of component 20 in ‘conflict’-related brain processes.

21) *21st Component*

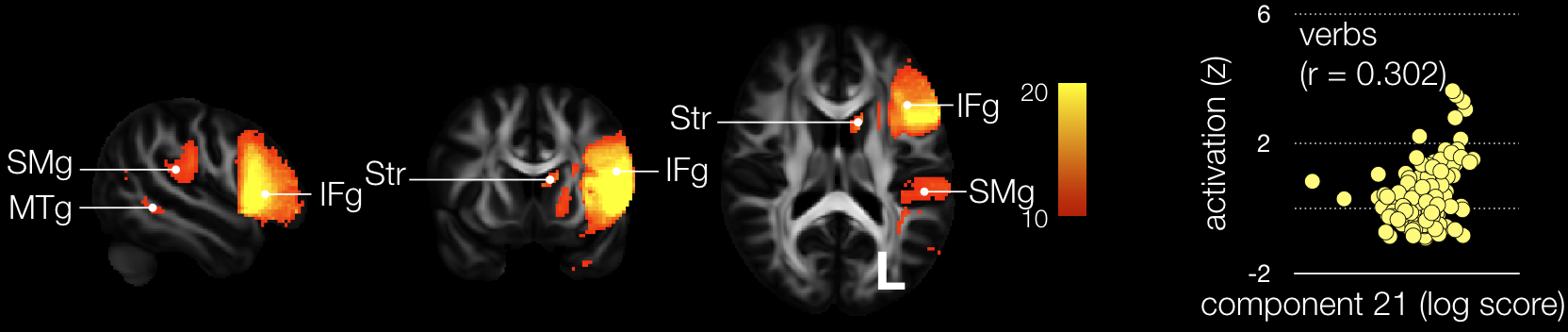

Supplementary figure 22: Twenty-first component map side-by-side with its correlation with task related functional MRI. SMg: Supramarginal gyrus, MTg: Middle Temporal gyrus, IFg: Inferior frontal gyrus, Str: Striatum.

Component 21 correlated significantly with ‘verbs’ related task-*f*MRI activations (r = 0.302).

Component 21 map was well replicable (r = 0.886) and significantly disconnected the inferior frontal gyrus, the supramarginal gyrus the middle temporal gyrus and the striatum in the left hemisphere. Verb generation has been reported as significantly impaired in patients with a stroke in the inferior frontal gyrus ^53^. Recent work also indicated a strong deficit in category fluency, which is a close clinical test to verb generation, after a disconnection of a functional network including the inferior frontal gyrus, part of the striatum, the supramarginal gyrus and the middle temporal gyrus ^54^. Thus previous report in stroke validate the potential involvement of component 21 in ‘verbs’ generation processes in the brain

22) *22nd Component*

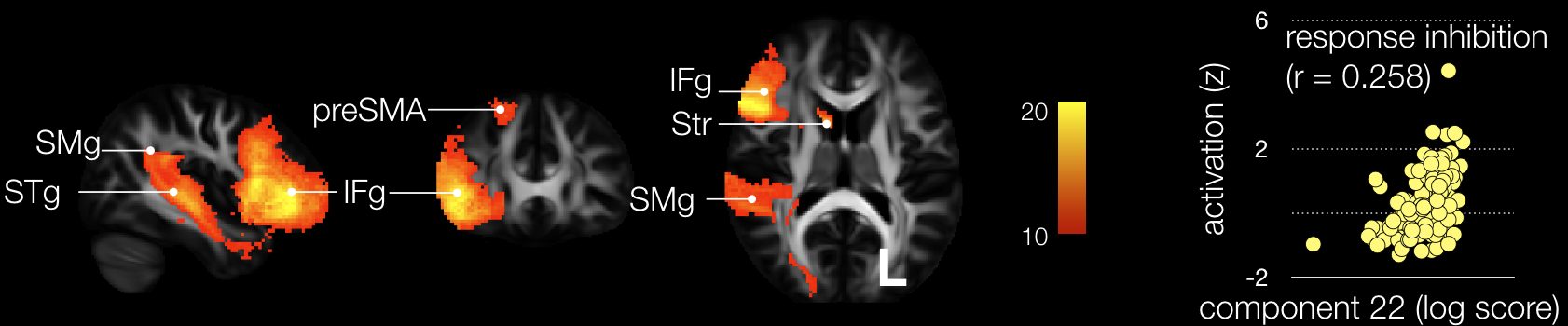

Supplementary figure 23: Twenty-second component map side-by-side with its correlation with task related functional MRI. SMg: Supramarginal gyrus, STg: Superior Temporal gyrus, IFg: Inferior frontal gyrus, preSMA: Presupplementary Motor Area, Str: Striatum.

Component 22 correlated significantly with ‘response inhibition’ related task-*f*MRI activations (r = 0.258). Component 22 map was moderately replicable (r = 0.750) and significantly disconnected the inferior frontal gyrus, the supramarginal gyrus, the superior temporal gyrus the presupplementary motor area and the striatum. Lesion to the inferior frontal gyrus is classically associated with response inhibition^55^, disorders in the inhibitory deficit during conflictual situations have been reported for lesions in the presupplementary area ^56^ and inhibition disorder are occurring after surgical resection of the inferior parietal lobule (i.e. including the supramarginal gyrus) ^57^. Lesion to the striatum also impairs inhibition ^58^, and specifically, in non-human primate, lead to the absence startle reflex inhibition ^59^. Hence lesion studies in human and non-human primates appears to support the involvement of component 22 in ‘response inhibition’ brain processes

23) *23rd Component*

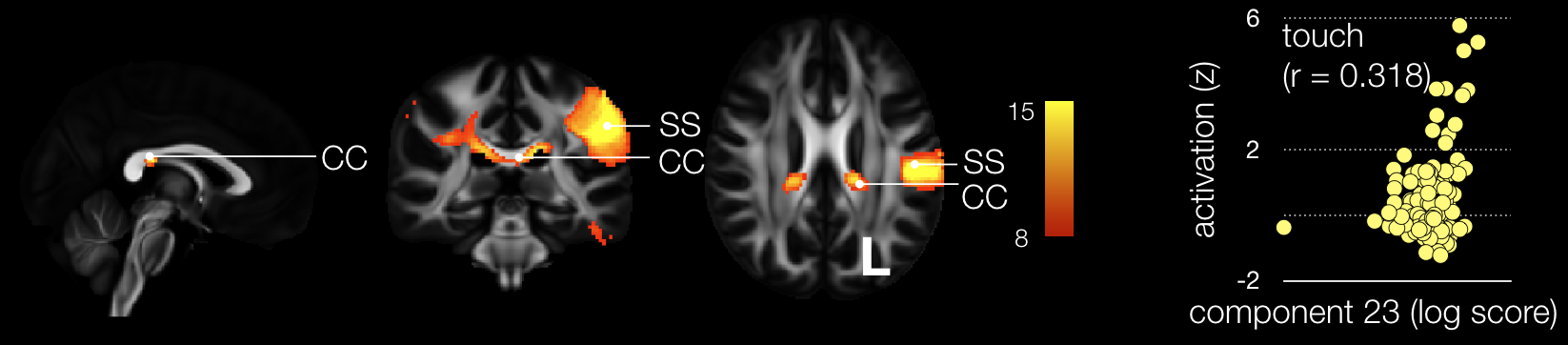

Supplementary figure 24: Twenty-third component map side-by-side with its correlation with task related functional MRI. SS: Somatosensory cortex, CC: Corpus Callosum.

Component 23 correlated significantly with ‘touch’ related task-*f*MRI activations (r = 0.318). Component 23 map was moderately replicable (r = 0.733) and significantly disconnected the left somatosensory cortex and the corpus callosum. Direct lesion to the somatosensory cortex or the disconnection of its subjacent white matter lead to numbness (i.e. somatosensory deficits) ^41^. However no lesion studies have, to our knowledge, reported somatosensory deficit after a disconnection of the corpus callosum. Hence the literature in lesion study only partially validate the implication of component 23 in ‘touch’ brain processes.

24) *24th Component*

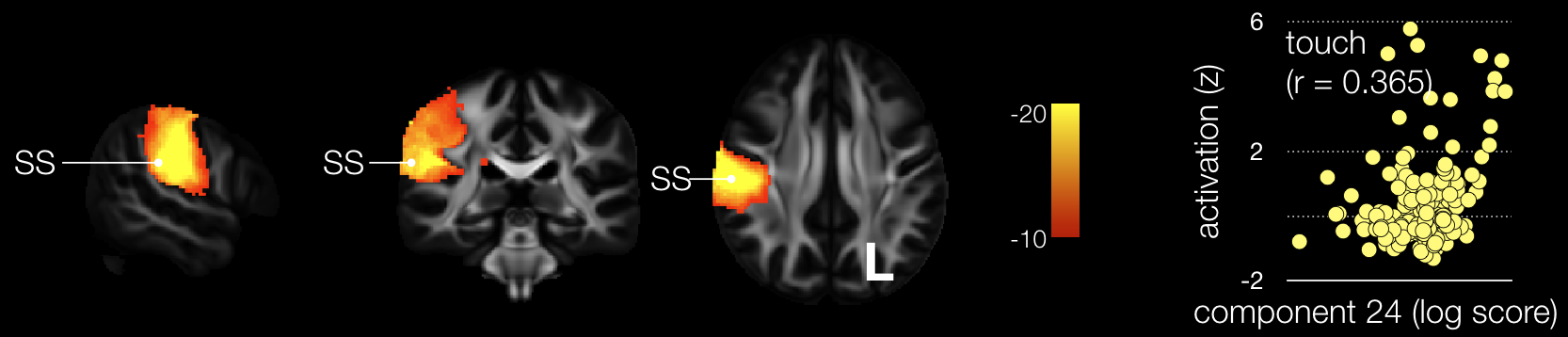

Supplementary figure 25: Twenty-fourth component map side-by-side with its correlation with task related functional MRI. SS: Somatosensory cortex

Component 24 correlated significantly with ‘touch’ related task-*f*MRI activations (r = 0.365). Component 24 map was moderately replicable (r = 0.718) and significantly disconnected the somatosensory cortex of the right hemisphere. As for component 23, direct lesion or disconnection of the somatosensory cortex is well known to lead to somatosensory deficits ^41^ and therefore validate the implication of component 24 in ‘touch’ related brain processes.

25) *25th Component*

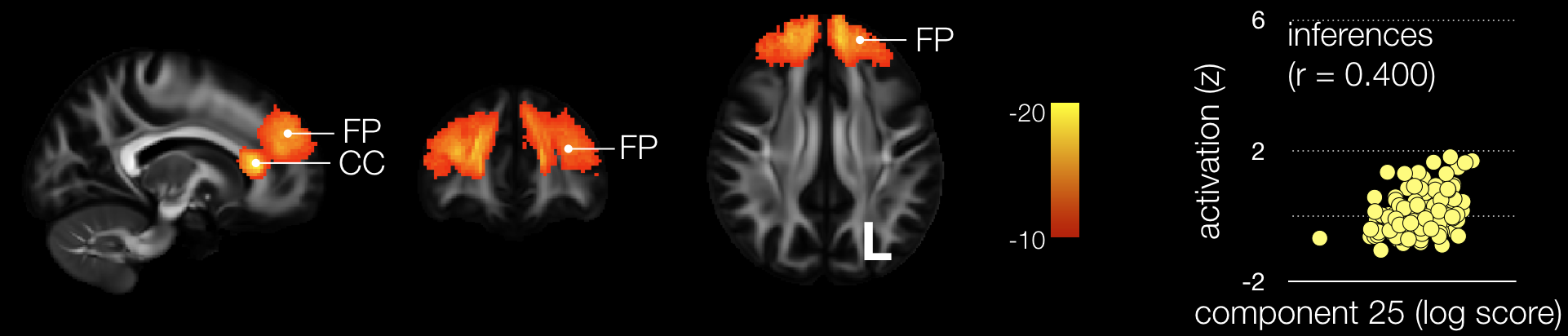

Supplementary figure 26: Twenty-fifth component map side-by-side with its correlation with task related functional MRI. CC: Corpus Callosum, FP: Frontal Pole

Component 25 correlated significantly with ‘inference’ related task-*f*MRI activations (r = 0.400). Component 25 map was well replicable (r = 0.854) and significantly disconnected the dorsal portion of the frontal pole together with its subjacent white matter including interhemispheric connections (i.e. corpus callosum)

Impairments in inferencing processes have been previously linked with right hemisphere damage (RHD) in the brain for many years ^60,61^. Early work suggested that adults with RHD were unable to generate inferences and comprehended only explicitly stated information ^62^ and the inferential deficit concerns specifically generation and integration ^63^. Despite some recent effort in further characterisation of this deficit, there is no article, to our knowledge, reporting a clear localisation of the lesions associated with deficits in inferential processes ^64^. However recent work in patients with brain lesions report an involvement of the left frontal pole and its connections for analogical reasoning, which require inference processes. The region reported would match the disconnection reported for component 25 in the left hemisphere and would partially validate the involvement of component 25 in ‘inference’ brain processes^65^.

26) *26th Component*

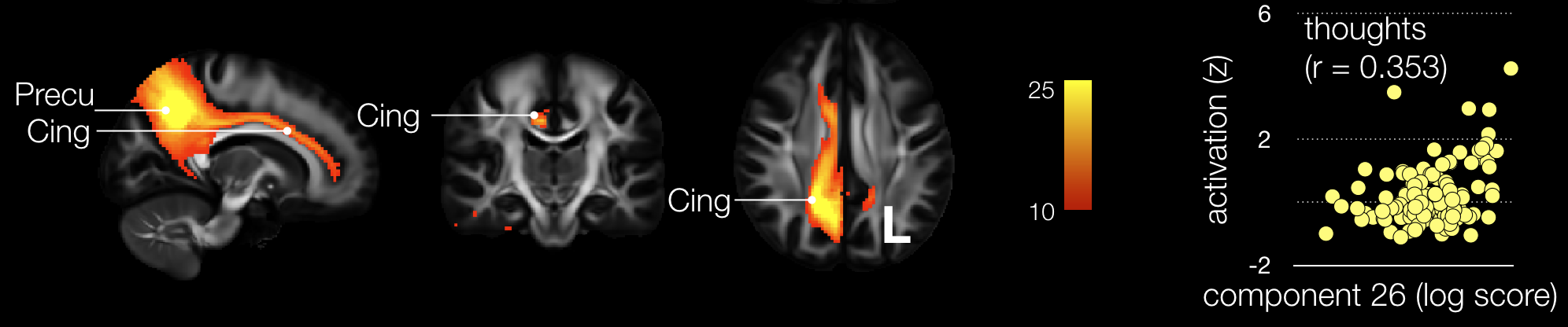

Supplementary figure 27: Twenty-sixth component map side-by-side with its correlation with task related functional MRI. Precu: Precuneus, Cing: Cingulum

Component 26 correlated significantly with ‘thought’ related task-*f*MRI activations (r = 0.353). Component 6 map was well replicable (r = 0.848) and significantly disconnected the dorsal portion of the cingulum together with the precuneus area. Brain lesions in these areas are uncommon for biological reasons. However, hypometabolism of the precuneus ^66^ as well as the direct electrical stimulation of the dorsal cingulum ^67^ frequently present with alteration of conscious state. Hence, since consciousness englobe introspection and thoughts^68^, the network associated with compenent 26 appears to be compatible with its involvement in ‘thought’ processes.

27) *27th Component*

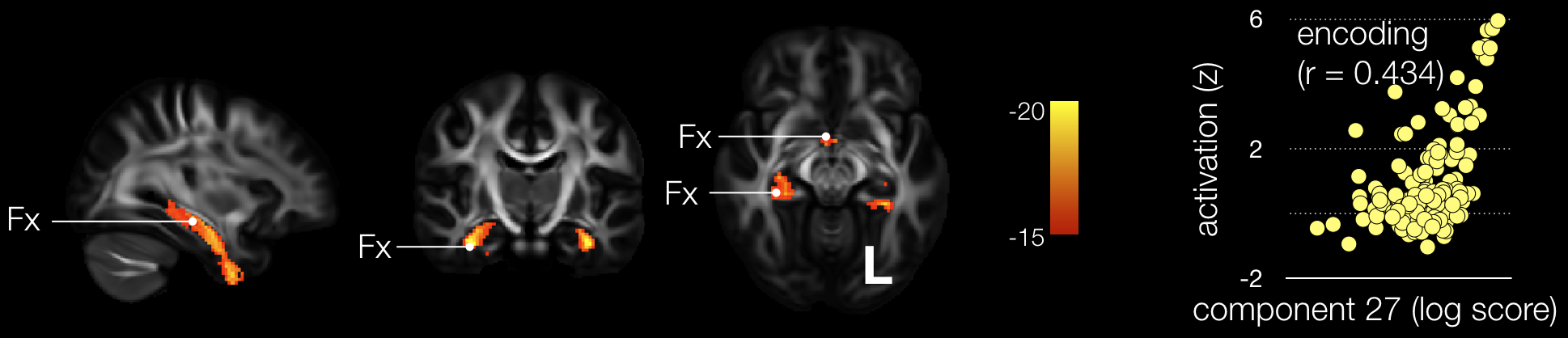

Supplementary figure 28: Twenty-seventh component map side-by-side with its correlation with task related functional MRI. Fx: Fornix

Component 27 correlated significantly with ‘encoding’ related task-*f*MRI activations (r = 0.353). Component 27 map was well replicable (r = 0.823) and significantly disconnected the fornix bilaterally. The fornix through its direct connection with the mamillary body and the hippocampus has a strong role in the encoding of episodic memories. Strokes in the fornix can occur and is systematically associated with anterograde amnesia ^69-74^. Additionally, the removal of a colloid cyst of the third ventricle during neurosurgery can damage the fornix and result in an anterograde amnesia ^75^. These results in neurologically impaired patients confirm the involvement of component 27 in ‘encoding’ processes.

28) *28th Component*

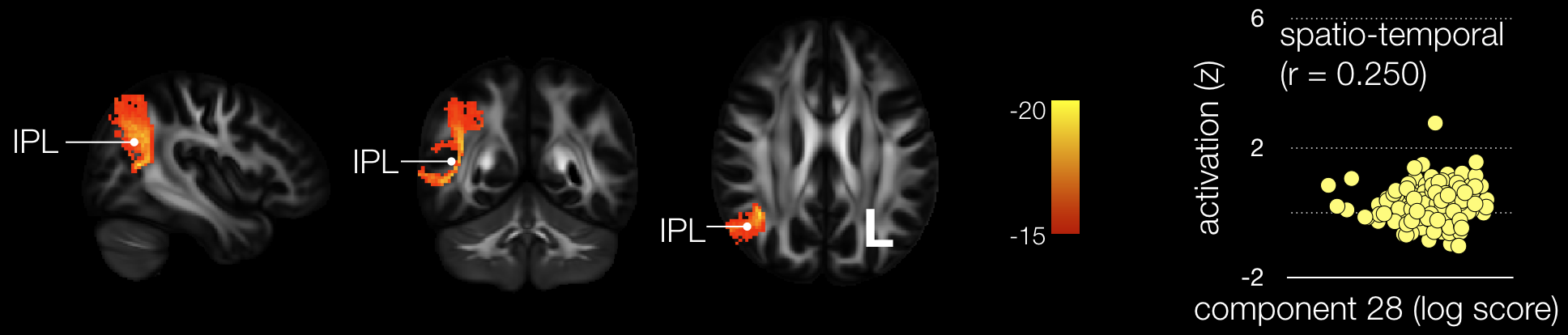

Supplementary figure 29: Twenty-eighth component map side-by-side with its correlation with task related functional MRI. IPL: Inferior Parietal Lobule

Component 28 correlated significantly with ‘spatio-temporal’ related task-*f*MRI activations (r = 0.250). Component 28 map was well replicable (r = 0.815) and significantly disconnected the right inferior parietal lobule and its subjacent short white matter interconnections. The right parietal lobe is well know for its spatial involvement but its functions extend beyond space to include control and sustainability of attention over time as well as detection of salient events ^76^ and representation of time ^77^. While the lesion of the parietal cortex and its connection has been well documented in term of hemispatial neglect ^39,78,79^, time perception deficits (e.g. dyschronometria) in stroke are poorly documented outside of cerebellar damages with only a single case report indicating the presence of signs of dyschronometria in a patient with a right parietal stroke ^80^. Hence, the involvement of component 28 in ‘spatio-temporal’ brain processes can be partially cross-

validated by the clinical signs reported in lesion studies.

29) *29th Component*

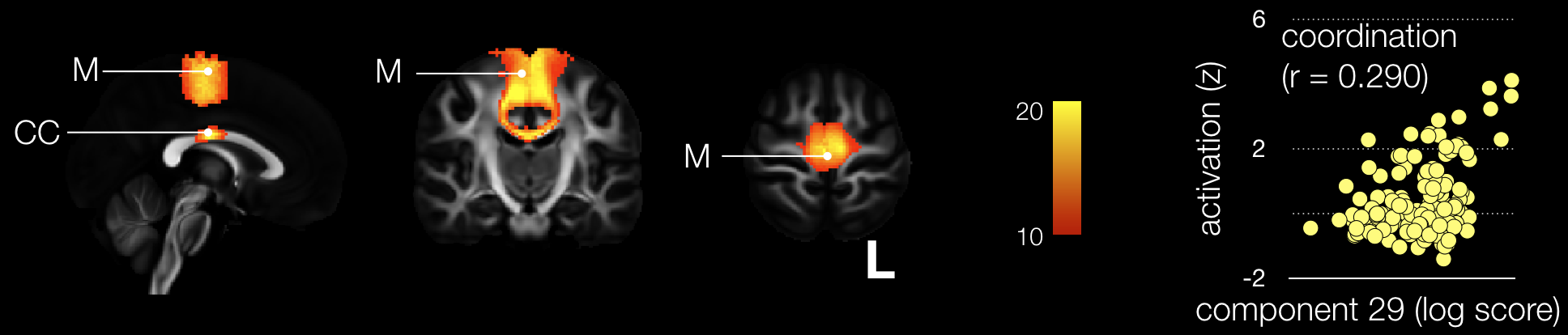

Supplementary figure 30: Twenty-ninth component map side-by-side with its correlation with task related functional MRI. M: primary motor cortex, CC: corpus callosum

Component 29 correlated significantly with ‘coordination’ related task-*f*MRI activations (r = 0.290). Component 29 map was moderately replicable (r = 0.769) and significantly disconnected the medial portion of the primary motor cortex together with its interhemispheric callosal connections. Lack of limb coordination has been reported after surgical disconnection of the anterior ^81^ and posterior portion of corpus callosum ^82^. Hence disconnections of the corpus callosum confirms the involvement of component 29 is ‘coordination’ brain processes.

30) *30th Component*

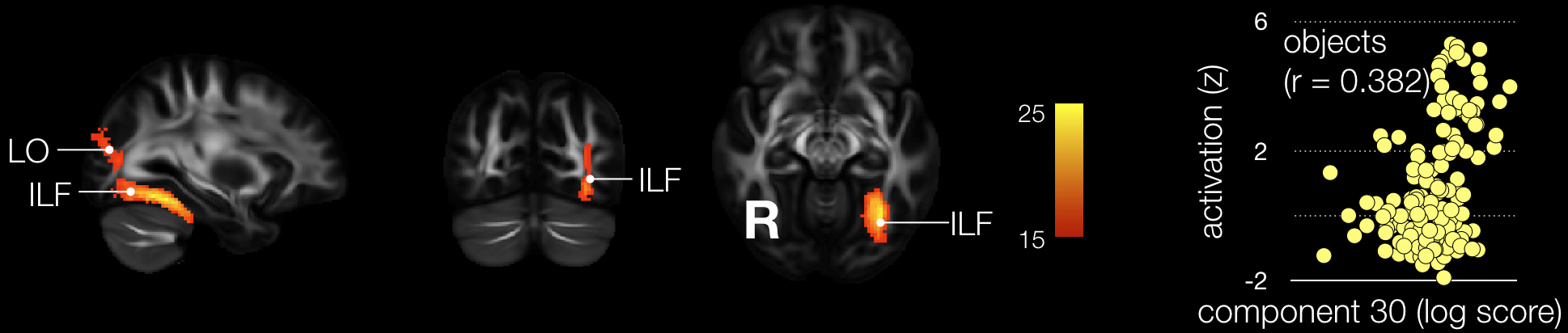

Supplementary figure 31: Thirtieth component map side-by-side with its correlation with task related functional MRI. LO: Lateral occipital, ILF: Inferior Longitudinal Fasciculus

Component 30 correlated significantly with ‘object’ related task-*f*MRI activations (r = 0.382)

Component 30 map was well replicable (r = 0.817) and significantly disconnected the lateral occipital region (i.e. LOP/LOC) and its subjacent white matter including the inferior longitudinal fasciculus in the left hemisphere. Stroke in the lateral occipital region lead to the incapacity to recognise objects either due the inability to group component parts of an object into a coherent whole (i.e., integrative agnosia)^83-85^ or to an alteration of early-level perceptual processing (i.e., apperceptive agnosia)^86,87^. Additionally visual agnosia has also been associated with the disconnection of the inferior longitudinal fasciculus^88^. Thus, previous research in lesioned brain appears to support the association of component 30 with ‘object’ related brain processes.

31) *31st Component*

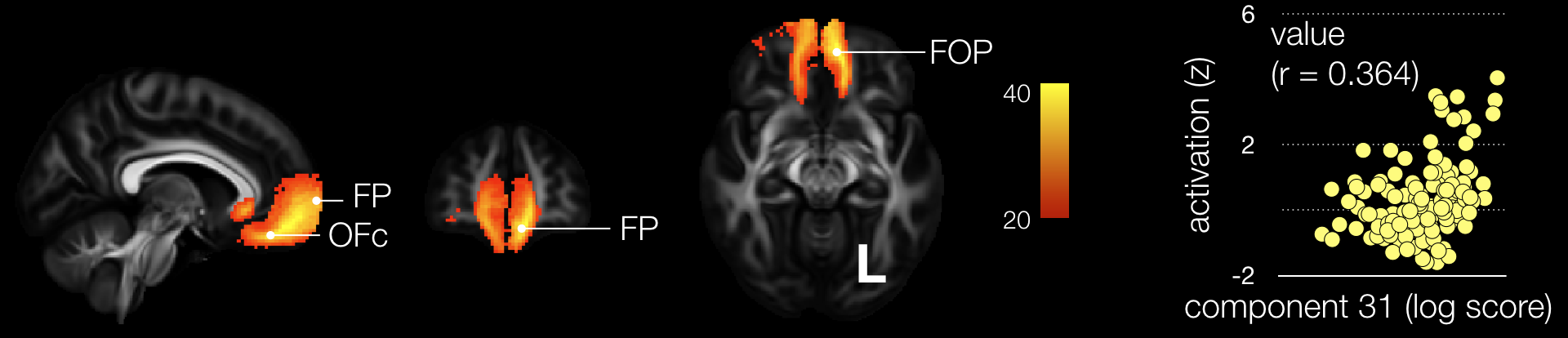

Supplementary figure 32: Thirty-first component map side-by-side with its correlation with task related functional MRI. OFc: Orbitofrontal cortex, FP: Frontal Pole, FOP: Frontal Orbito Polar tract.

Component 31 correlated significantly with ‘value’ related task-*f*MRI activations (r = 0.364)

Component 31 map was highly replicable (r = 0.907) and significantly disconnected the orbito-frontal cortex bilaterally and its interhemispheric connections (i.e. corpus callosum).

Subjective value representation has been documented after a lesion in the orbitofrontal cortex in the case of Phineas Gage who did ‘*not estimate size or money acutely, though he has memory as perfect as ever. He would not take $1000 for a few pebbles which he took from an ancient river bed where he was at work*’ ^89^. Note that Phineas Gage brain damage also disconnected the orbitofrontal cortex from the frontal pole ^90^ (i.e. frontal orbito-polar tract ^42,91^). Direct lesion of the orbitofrontal cortex is also classically linked with the impairment of subjective value representation in non-human primates. To date, there is no study, to our knowledge, reporting an involvement of interhemispheric connections in the processing or representation of values. Therefore, classical studies of the orbitofrontal cortex are partially supportive of the involvement of component 31 in ‘values’ brain processes.

32) *32nd Component*

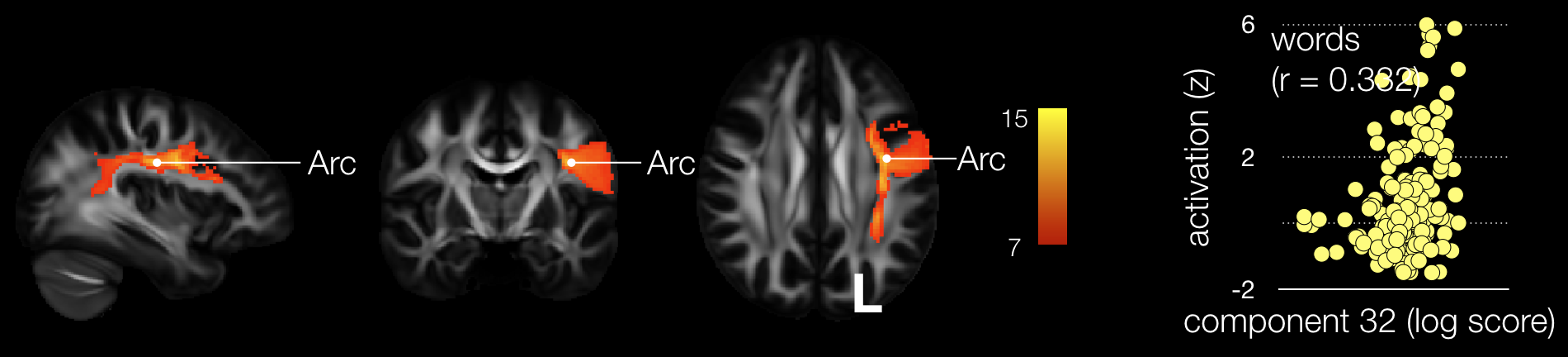

Supplementary figure 33: Thirty-second component map side-by-side with its correlation with task related functional MRI. Arc: Arcuate fasciculus

Component 32 correlated significantly with ‘words’ related task-*f*MRI activations (r = 0.332)

Component 32 map was moderately replicable (r = 0.711) and significantly disconnected the arcuate fasciculus. The role of the arcuate fasciculus in the oral production of words has been revisited in the original case of ‘Tan’ ^90,92^ but also in group studies ^3^. Lesion to the arcuate fasciculus has also been reported with deficits in words comprehension^93^. All in all studies of brain lesions support the involvement of component 32 in ‘words’ brain processing.

33) *33rd Component*

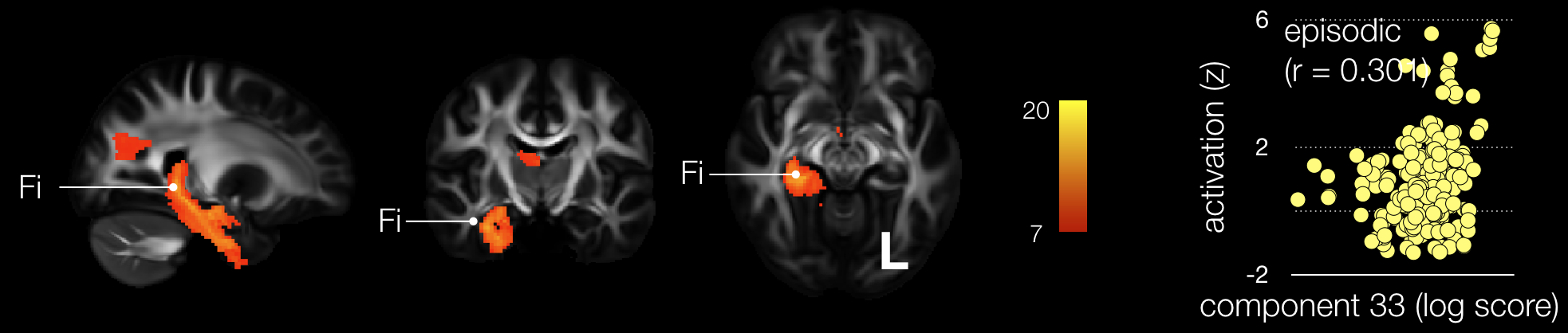

Supplementary figure 34: Thirty-third component map side-by-side with its correlation with task related functional MRI. Fi: Fimbria

Component 33 correlated significantly with ‘episodic’ related task-*f*MRI activations (r = 0.301). Component 33 map was poorly replicable (r = 0.669) and significantly disconnected the posterior-inferior part of the Fornix (i.e. the fimbria). The fimbria emerges from the hippocampus in the medial temporal lobe and project into the body of the fornix medially. Disconnection of the medial temporal lobe after a stroke has been significantly associated with impaired episodic memory ^94^. Additionally the case of H.M. critically documented a severe anterograde amnesia after a bilateral damage of the medial temporal lobe that included a disconnection of the fimbria ^90^. Hence the involvement of component 33 in ‘episodic’ brain processes appears to be supported by studies in lesioned brains.

34) *34th Component*

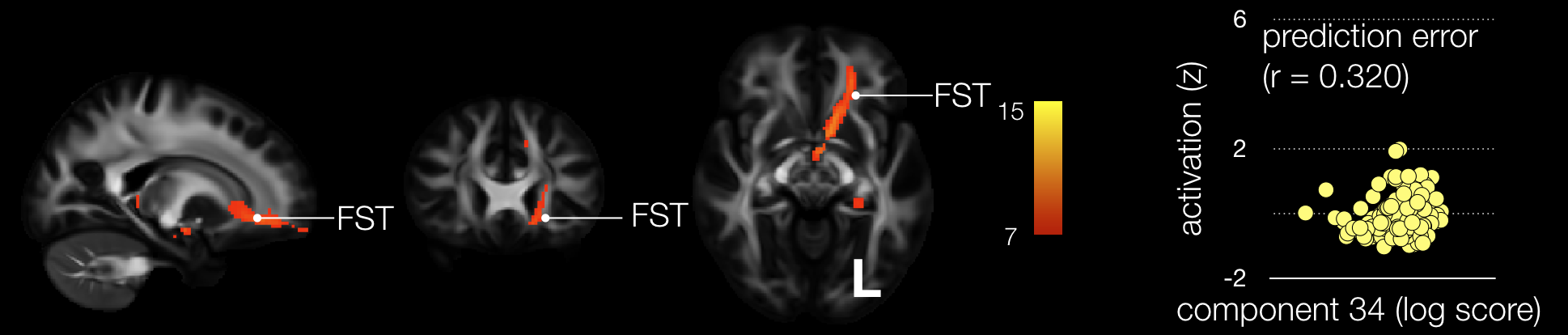

Supplementary figure 35: Thirty-forth component map side-by-side with its correlation with task related functional MRI. FST: Fronto-striatal tract

Component 34 correlated significantly with ‘prediction error’ related task-*f*MRI activations (r = 0.301). Component 34 map was poorly replicable (r = 0.393) and significantly disconnected the fronto-striatal tract. Prediction error together with cognitive flexibility can typically be assessed in patient with stroke using the Wisconsin Card Sorting test ^95^. Lesions leading to perseverative errors were located in the white matter of the prefrontal cortex encompassing the dorsolateral prefrontal cortex and the frontal pole, stretching to the posterior frontal and parietal cortex ^96^. This pattern of lesion is compatible with the involvement of the component 34 in ‘prediction error’ brain processes.

35) *35th Component*

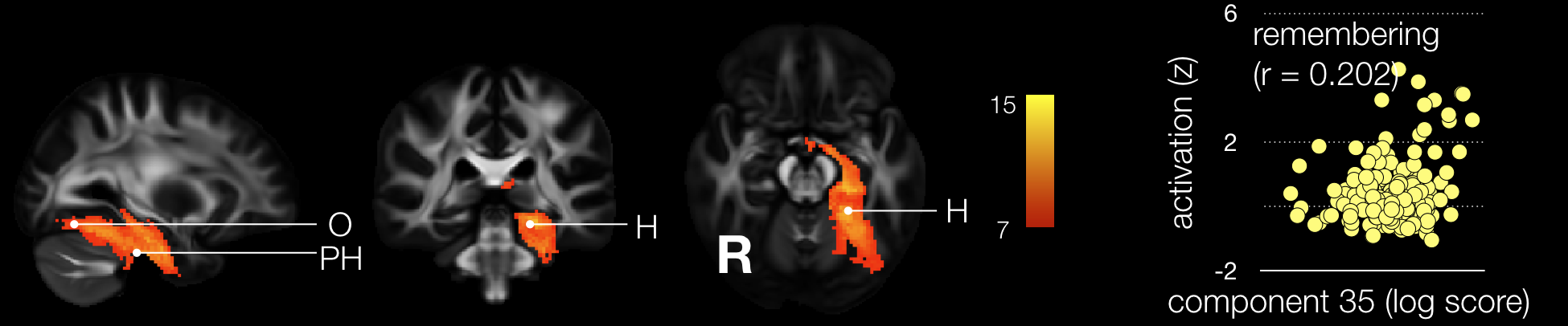

Supplementary figure 36: Thirty-fifth component map side-by-side with its correlation with task related functional MRI. FST: Fronto-striatal tract

Component 35 correlated significantly with ‘remembering’ related task-*f*MRI activations (r = 0.301). Component 6 map was poorly replicable (r = 0.660) and significantly disconnected the occipital lobe from the hippocampus and the medial temporal lobe in the left hemisphere. A single case patient with a lesion in the occipital lobe have been reported with an implicit memory deficit dissociated from his explicit performance ^97^. Another report indicate that patients with posterior cerebral infarction involving the white matter of the occipital lobe connecting the hippocampus and the parahippocampus gyrus are impaired at drawing objects from memory ^98^. However, it is important to stress that all these cases had their lesion located in the right hemisphere. Hence, lesions studies appear to only partially support the involvement of component 35 in ‘remembering’ brain processes.

36) *36th Component*

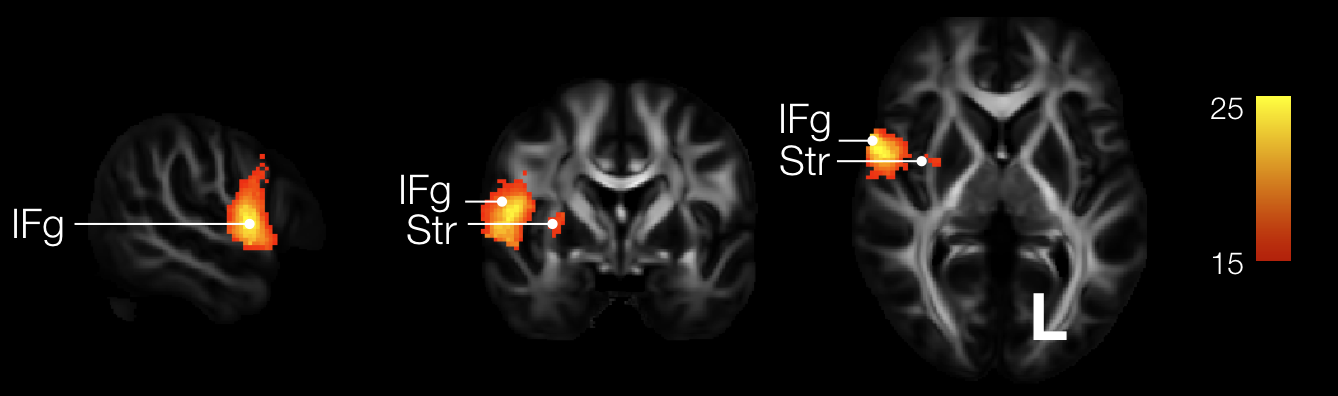

Supplementary figure 37: Thirty-sixth component map. IFg: Inferior Frontal gyrus, Str: Striatum.

Component 36 map was moderately replicable (r = 0.768) and significantly disconnected the posterior part of the inferior gyrus (i.e. pars opercularis) and the striatum in the right hemisphere. Correlations between component 36 and task-related functional activation did not survive Bonferroni correction for multiple comparisons (n = 590 functions).

37) *37th Component*

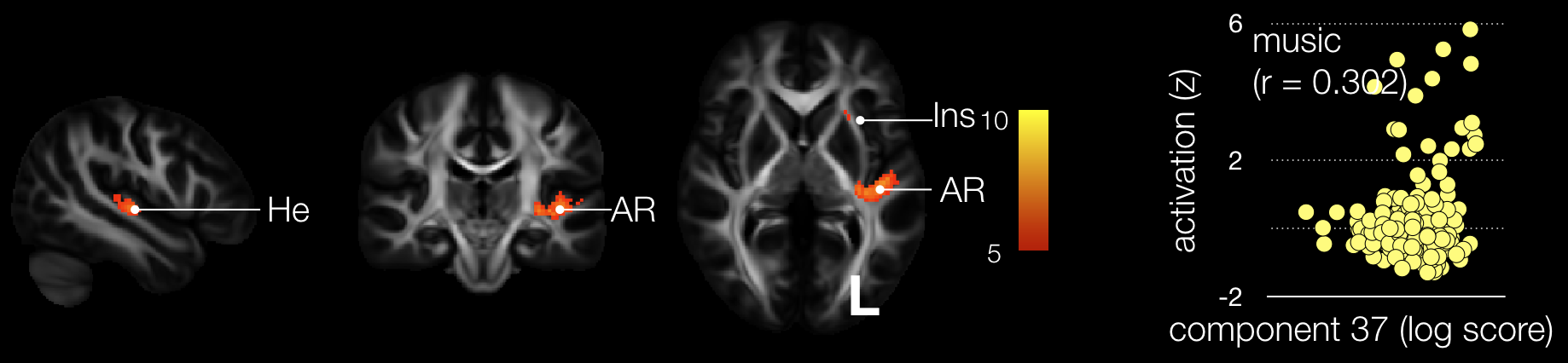

Supplementary figure 38: Thirty-seventh component map side-by-side with its correlation with task related functional MRI. He: Heschl’s gyrus, AR: acoustic radiations, Ins: Insula

Component 37 correlated significantly with ‘music’ related task-*f*MRI activations (r = 0.301).

Component 6 map was well replicable (r = 0.807) and significantly disconnected the Heschl’s gyrus, the acoustic radiations^99^ and the insula in the left hemisphere. Amusia acquired after a stroke in the right hemisphere has previously been reported for lesions involving the temporal gyrus and the insula ^100,101^. Since amusia have been reported for both left and right brain damage ^102^, these results confirms the involvement of component 37 in ‘music’ brain processes.

38) *38th Component*

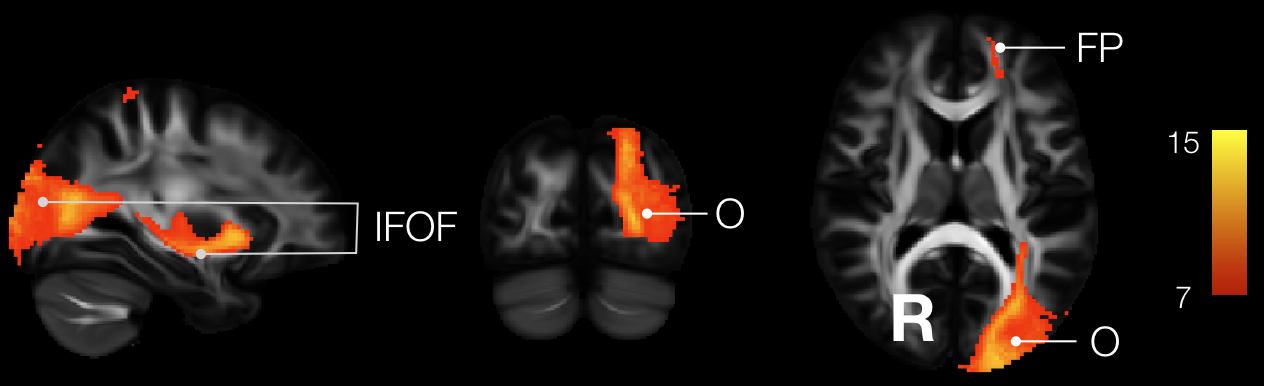

Supplementary figure 39: Thirty-eighth component map. IFOF: Inferior Fronto-Occipital Fasciculus, O: Occipital lobe, FP: Frontal Pole.

Component 38 map was moderately replicable (r = 0.771) and significantly disconnected a portion of the inferior fronto-occipital fasciculus that connects the occipital lobe to the frontal pole in the left hemisphere. Correlations between component 38 and task-related functional activation did not survive Bonferroni correction for multiple comparisons (n = 590 functions).

39) *39th Component*

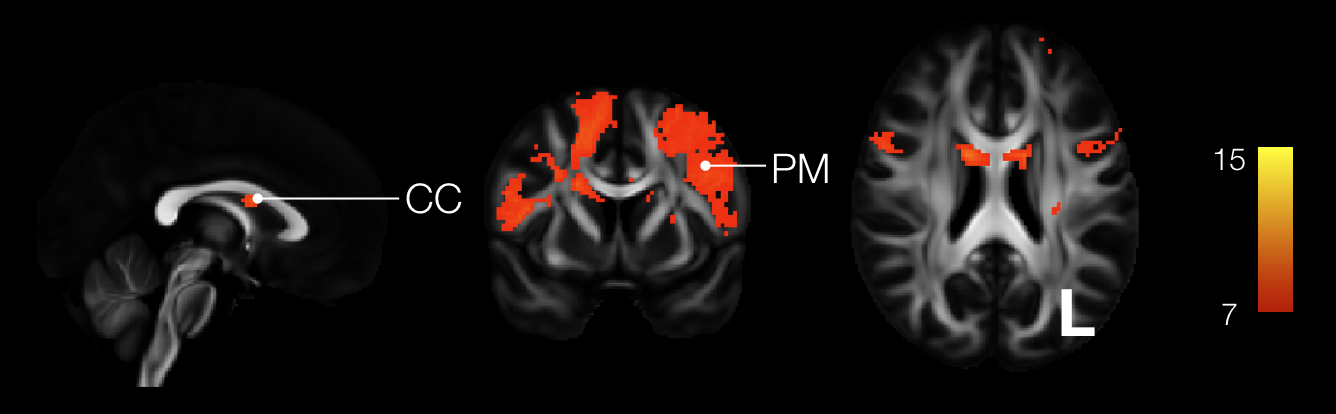

Supplementary figure 40: Thirty-ninth component map. PM: premotor cortex, CC: Corpus callosum

Component 39 map was poorly replicable (r = 0.569) and significantly disconnected the premotor cortex in both hemispheres as well as their interhemispheric connections. Correlations between component 39 and task-related functional activation did not survive Bonferroni correction for multiple comparisons (n = 590 functions).

40) *40th Component*

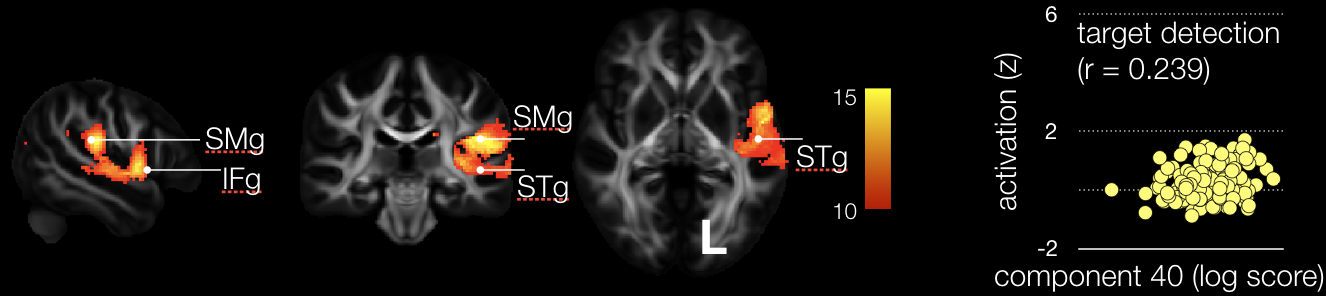

Supplementary figure 41: Fortieth component map with its correlation with task related functional MRI. SMg: SupraMarginal gyrus, IFg: Inferior Frontal gyrus, STg: Superior Temporal gyrus

Component 40 correlated significantly with ‘target detection’ related task-*f*MRI activations (r = 0.301). Component 40 map was moderately replicable (r = 0.729) and significantly disconnected the posterior part of the inferior frontal gyrus, the supramarginal gyrus and the superior temporal gyrus as well as their subjacent white matter in the left hemisphere. The areas are classically involved in visual neglect in the right hemisphere^103,104^. Considering the occurrence of visual neglect after a stroke in the left hemisphere lesion^105^ we can assume that the literature in brain lesion support the involvement of component 40 in ‘target detection’ processes.

41) *41st Component*

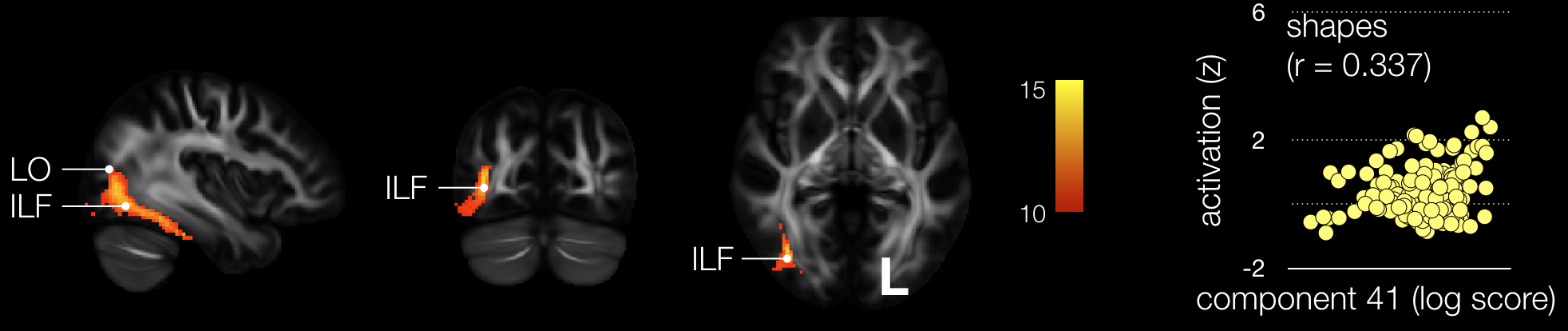

Supplementary figure 42: Forty-first component map with its correlation with task related functional MRI. LO: Lateral occipital, ILF: Inferior Longitudinal Fasciculus

Component 41 correlated significantly with ‘target detection’ related task-*f*MRI activations (r = 0.337). Component 41 map mirrored component 30 was poorly replicable (r = 0.696) and significantly disconnected the lateral occipital region (i.e. LOP/LOC) and its subjacent white matter including the inferior longitudinal fasciculus in the right hemisphere. As for component 30, stroke in the lateral occipital region lead to the incapacity to recognise objects either due the inability to group component parts of an object into a coherent shape (i.e., integrative agnosia)^83-85^ or to an alteration of early-level perceptual processing (i.e., apperceptive agnosia)^86,87^. Additionally visual agnosia has also been associated with the disconnection of the inferior longitudinal fasciculus^88^. Thus, previous research in lesioned brain appears to support the association of component 41 with ‘shape’ related brain processes.

42) *42nd Component*

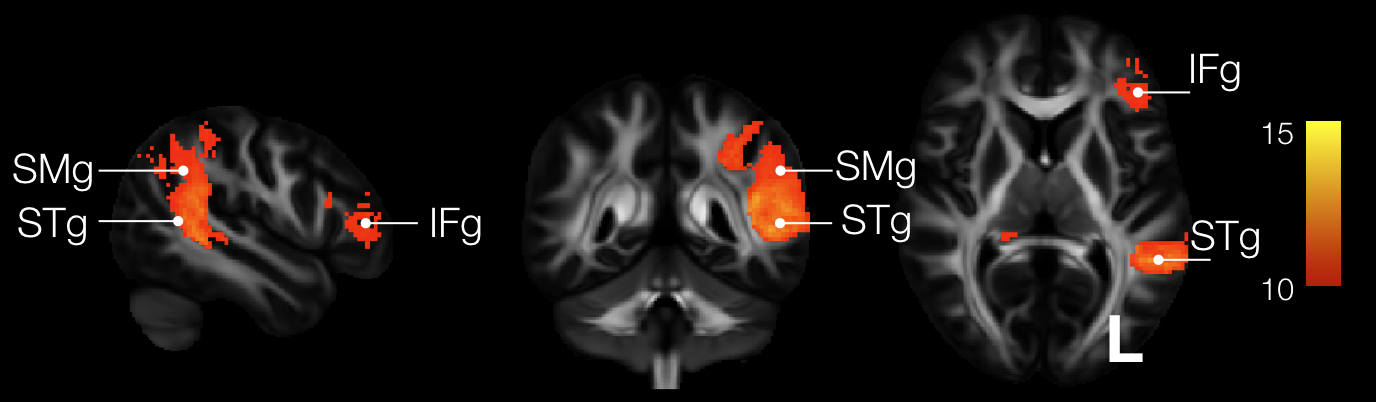

Supplementary figure 43: Forty-second component map with its correlation with task related functional MRI. SMg: SupraMarginal gyrus, IFg: Inferior Frontal gyrus, STg: Superior Temporal gyrus.

Component 42 map was moderately replicable (r = 0.770) and significantly disconnected the supramarginal and the superior temporal gyri as well in the mid portion of the inferior frontal gyrus (i.e. pars triangularis). Correlations between component 42 and task-related functional activation did not survive Bonferroni correction for multiple comparisons (n = 590 functions).

43) *43rd Component*

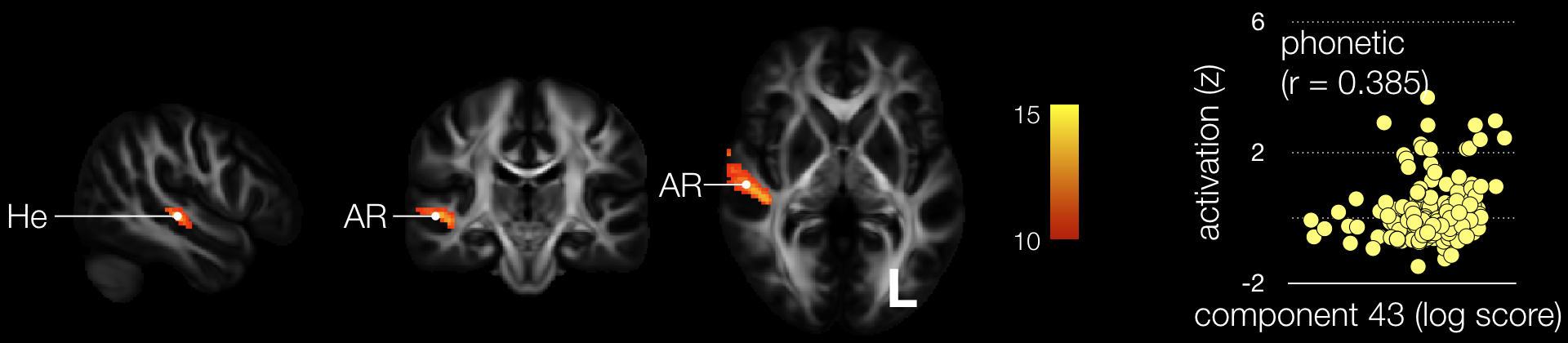

Supplementary figure 44: Forty-third component map with its correlation with task related functional MRI. He: Heschl’s gyrus, AR: acoustic radiations

Component 43 correlated significantly with ‘phonetic’ related task-*f*MRI activations (r = 0.385). Component 43 map was poorly replicable (r = 0.540) and significantly disconnected the Heschl gyrus and its subjacent white matter connections including the acoustic radiations^99^ in the right hemisphere. Stroke are involving the auditory system produces psychoacoustic dysfunctions^106^. Thus, stroke lesion partially validate the involvement of component 43 in phonetic disorders.

44) *44th Component*

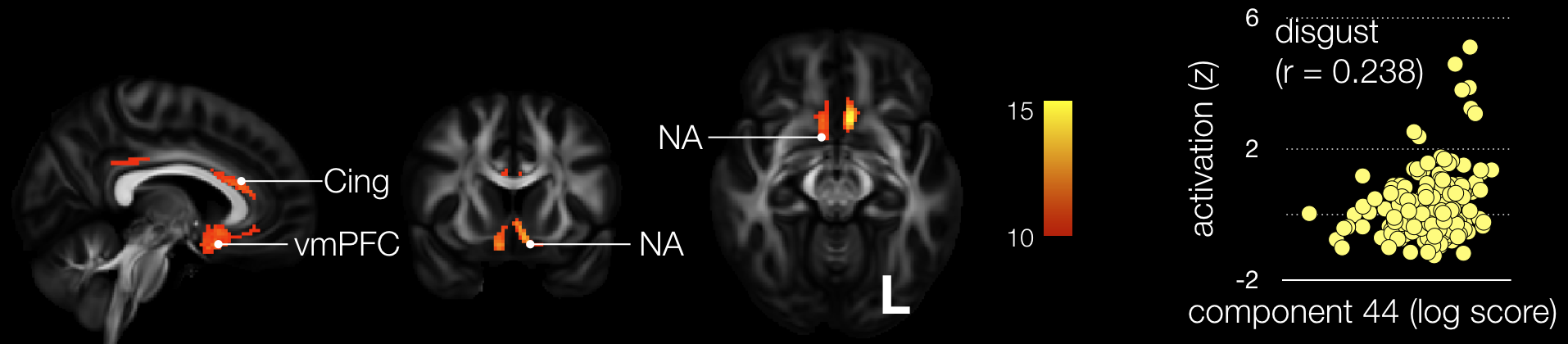

Supplementary figure 45: Forty-forth component map with its correlation with task related functional MRI. Cing: Cingulum, vmPFC: ventromedial PreFrontal Cortex, Acc: Accumbens nucleus

Component 44 correlated significantly with ‘disgust’ related task-*f*MRI activations (r = 0.385). Component 44 map was moderately replicable (r = 0.723) and significantly disconnected the anterior cingulate cortex, the ventromedial prefrontal cortex and the accumbens nucleus. Anterior cingulotomy lead patient to not be able to recognize negative expressions (i.e. fear, disgust, anger and sadness) while the recognition of positive and neutral expression remains unimpaired ^107^. Loss of interpersonal disgust can occur after a lesion to the ventromedial prefrontal cortex ^108^ in humans and pharmacological inhibition of the accumbens nucleus cause excessive disgust in rats ^109^.

45) *45th Component*

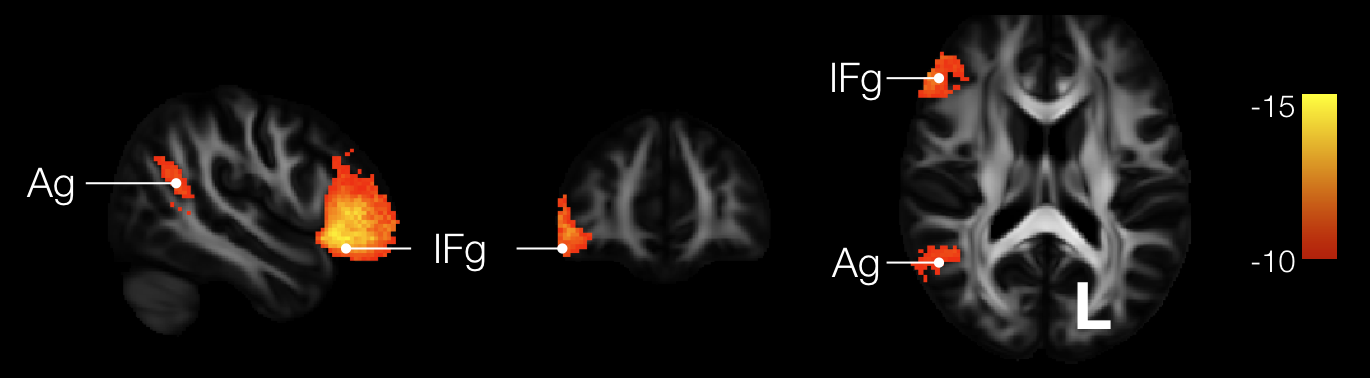

Supplementary figure 46: Forty-fifth component map. Ag: Angular gyrus, IFg: Inferior Frontal gyrus

Component 45 map was poorly replicable (r = 0.675) and significantly disconnected the angular gyrus from the inferior frontal gyrus (i.e. pars triangularis) in the right hemisphere. Correlations between component 45 and task-related functional activation did not survive Bonferroni correction for multiple comparisons (n = 590 functions).

46) *46th Component*

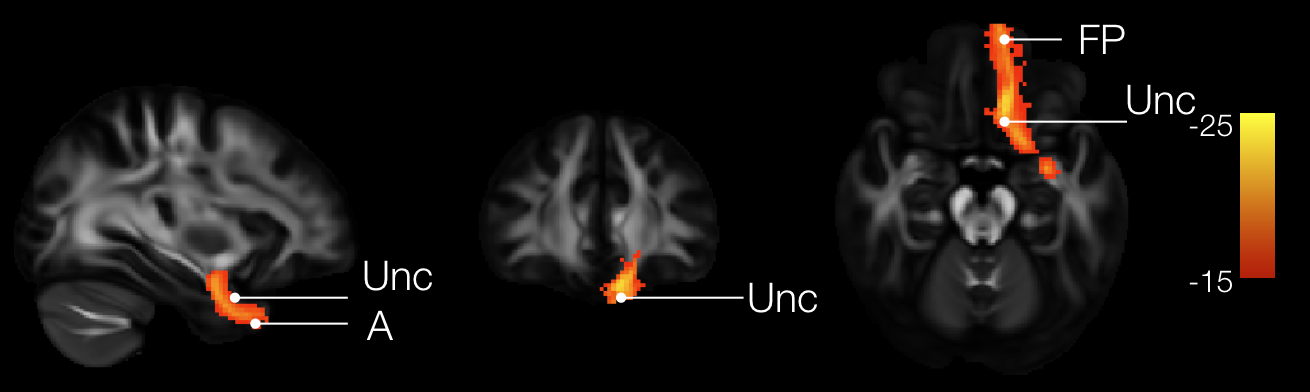

Supplementary figure 47: Forty-sixth component map. Unc: Uncinate fasciculus, A: Amygdala, FP: Frontal Pole

Component 6 map was well replicable (r = 0.870) and significantly disconnected a portion of the uncinate fasciculus connecting the amygdala to the frontal pole. Correlations between component 46 and task-related functional activation did not survive Bonferroni correction for multiple comparisons (n = 590 functions).

**B) Example of left and right finger tapping activation mapped onto the white matter**

Example of permuted linear regression applied to identify the statistical contribution of each voxel of each component map (i.e. independent variables) to real group-level fMRI maps (i.e. the dependent variable). The real group-level fMRI maps were derived from a left and a right finger tapping task in 142 right-handed participants ^110^.

The 46 disconnectome components succeeded at explaining 66% of the variance of the left finger tapping activation and 63% of the variance of the right finger tapping activation.

Since we have two sets of component maps, mapping of left and right finger tapping was mapped onto the white matter twice. Replicability of these mappings was assessed using a Pearson correlation between the results and indicated excellent replicability for the left (92%) and the right (94%) mapping onto the white matter.

Finger tapping activated the contralateral motor cortex, the premotor cortex and the supplementary motor area. The mapping of both activations onto the white matter indicated a significant involvement of the corticospinal tract, the corpus callosum, frontostriatal connections as well as additional (unlabeled) connection to, the motor and the premotor cortices.

Finally, two typical lesions showed disconnections sharing 44% of the variance the pattern of connection revealed by the finger-tapping mapped onto white matter. Hence these lesions are very likely to disconnect the white matter pathways ‘feeding’ into the pattern of activation for left and right finger tapping respectively putatively leading to motor symptoms of the left or the right hand.

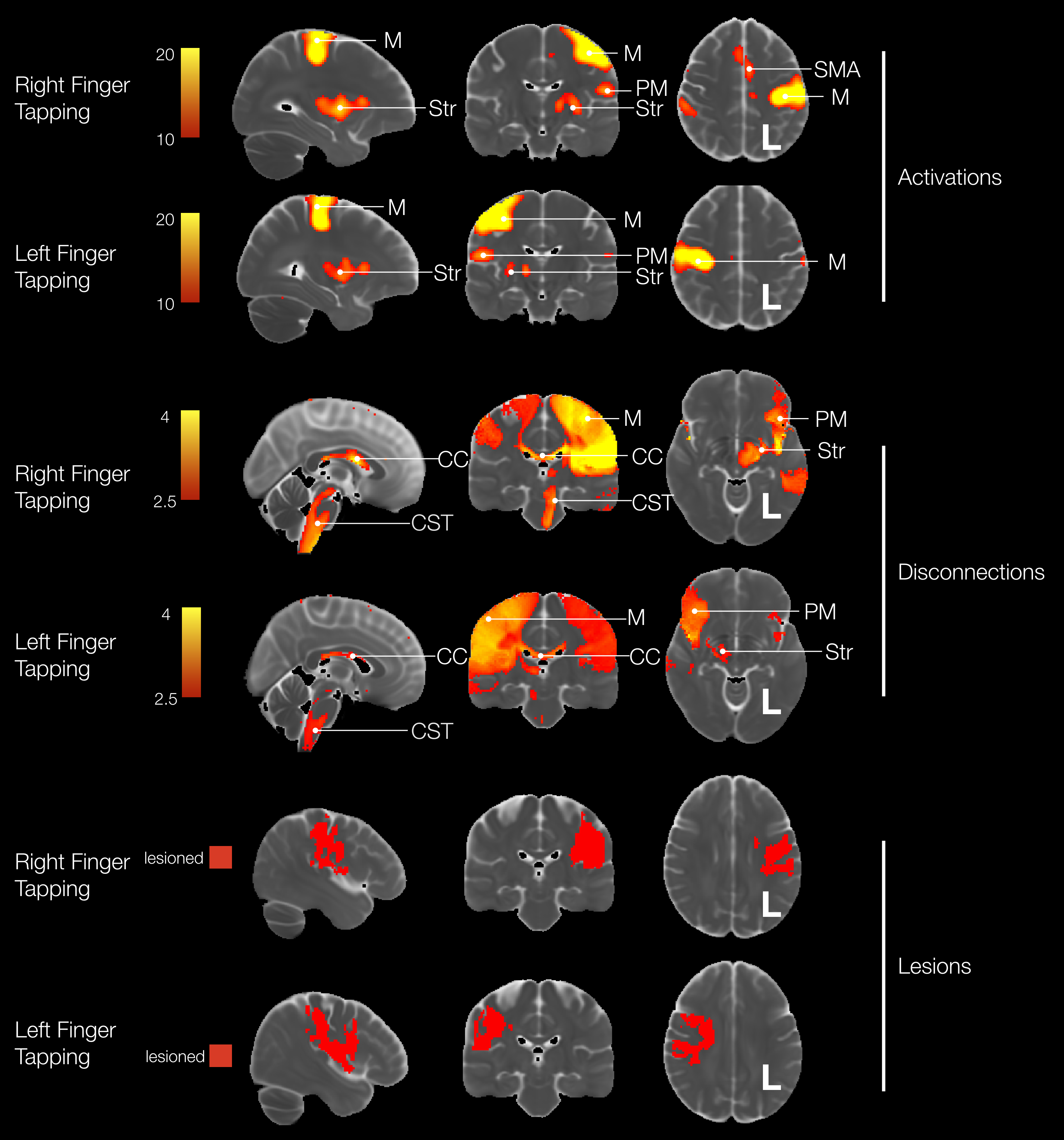

Supplementary figure 48: Mapping of activation onto the white matter and typical lesion leading to the disconnection of the network of activations. M: Motor cortex, PM: Premotor cortex, Str: Striatum, SMA: Supplementary Motor Area, CST: Cortico-Striatal Tract, CC: Corpus Callosum.

**C) High resolution of the atlas of white matter function**

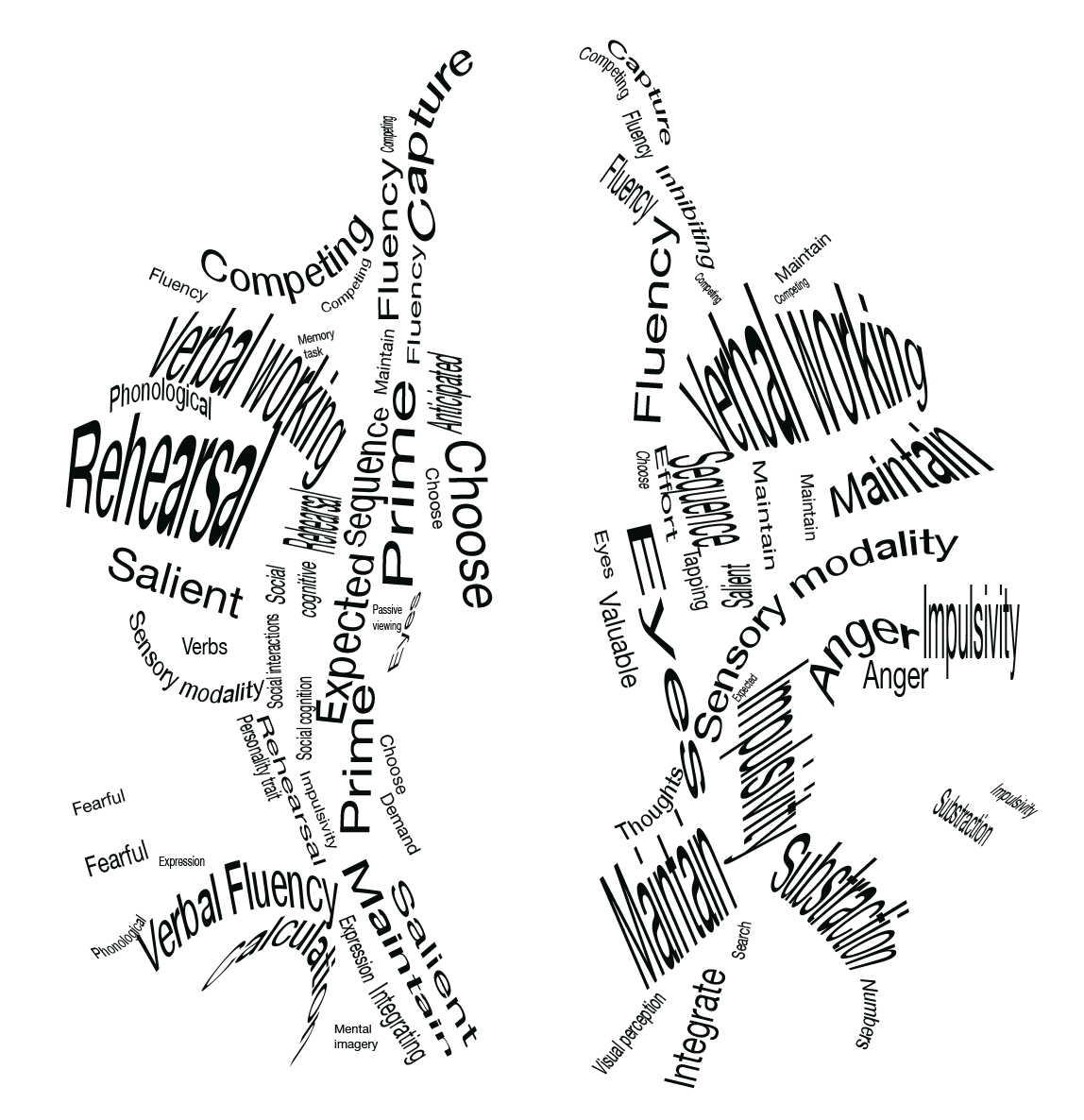

Supplementary figure 49: atlas of white matter function MNI slice z = 50 (neurological convention)

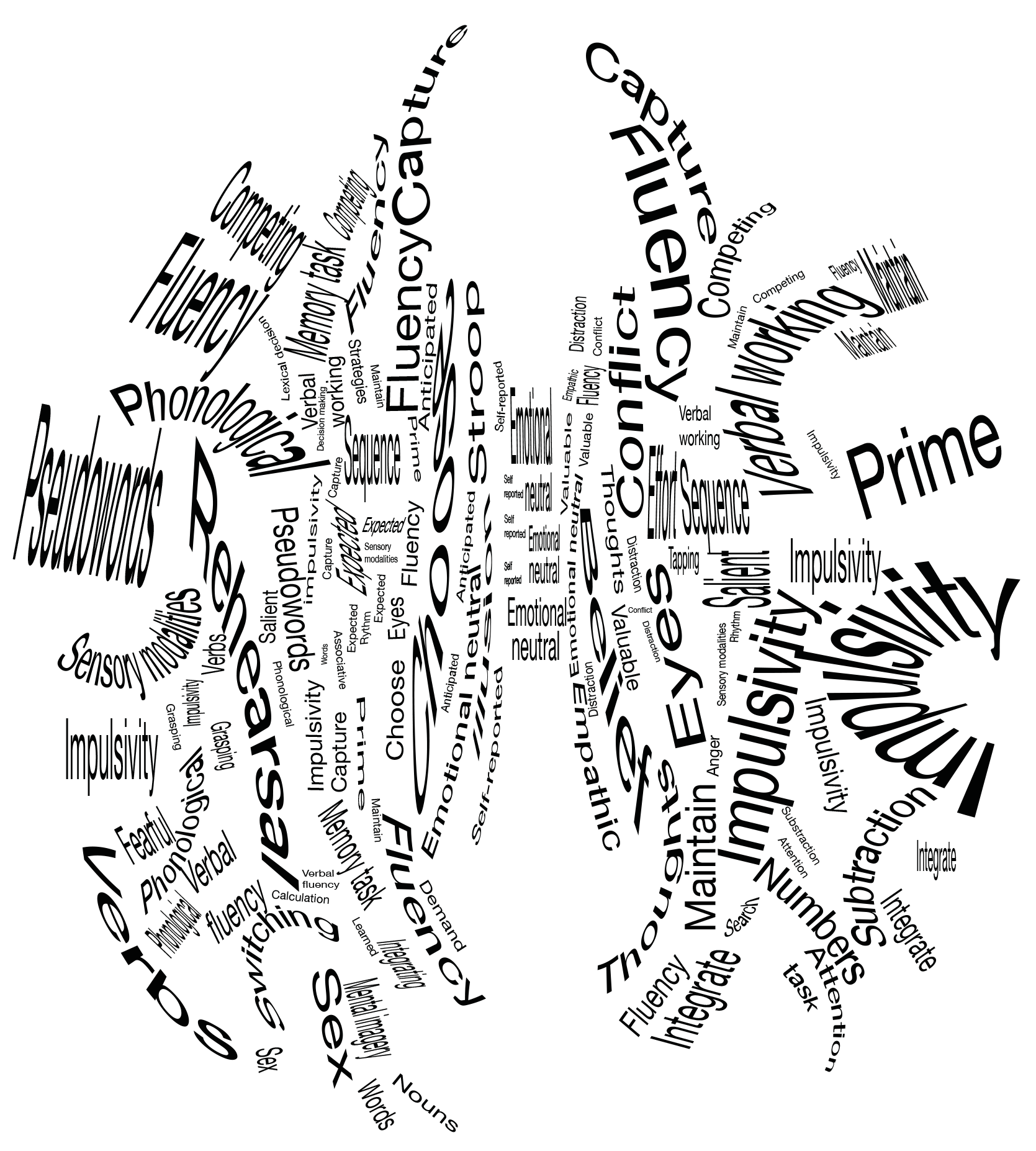

Supplementary figure 50: atlas of white matter function MNI slice z = 40 (neurological convention)

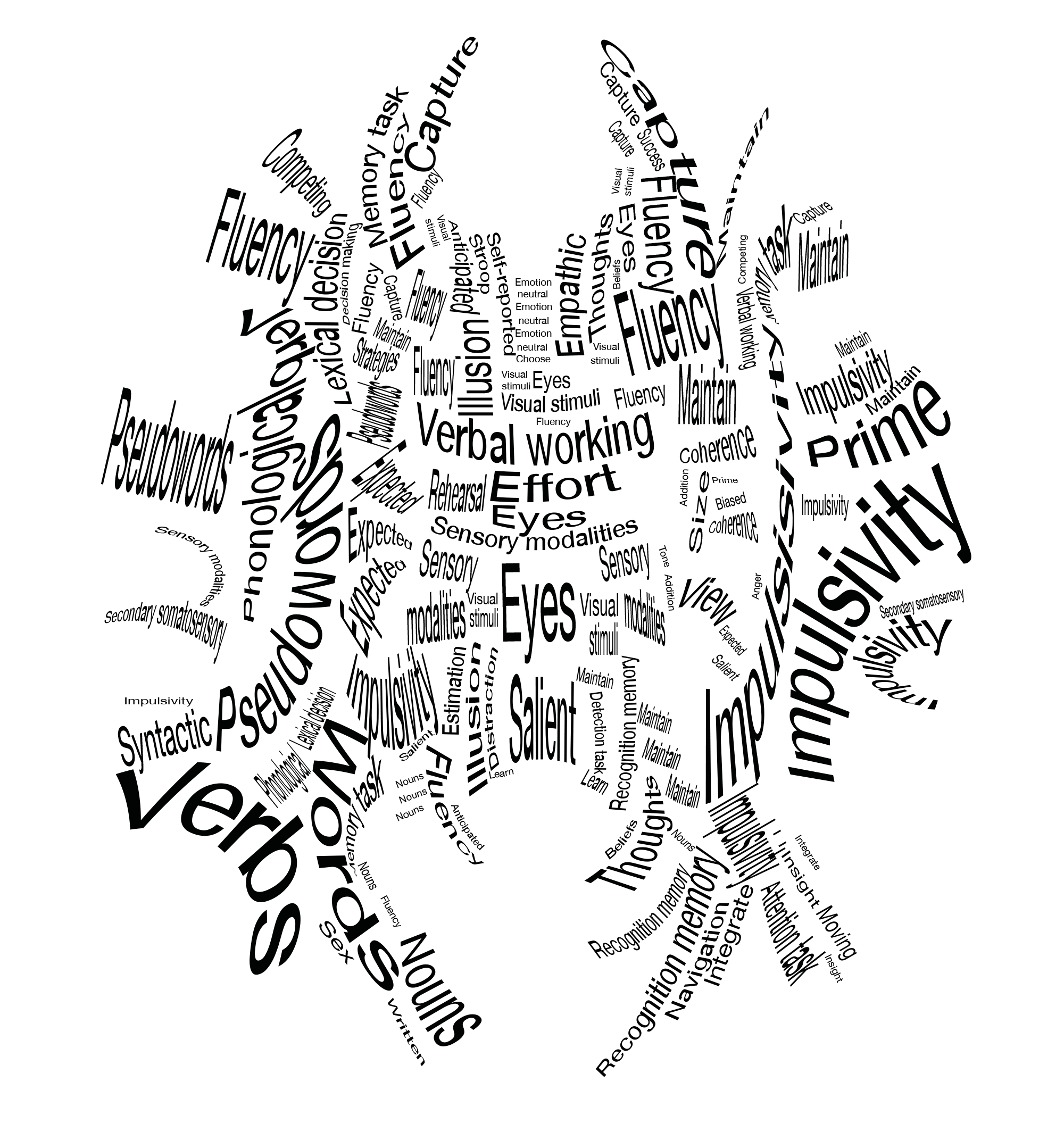

Supplementary figure 51: atlas of white matter function MNI slice z = 30 (neurological convention)

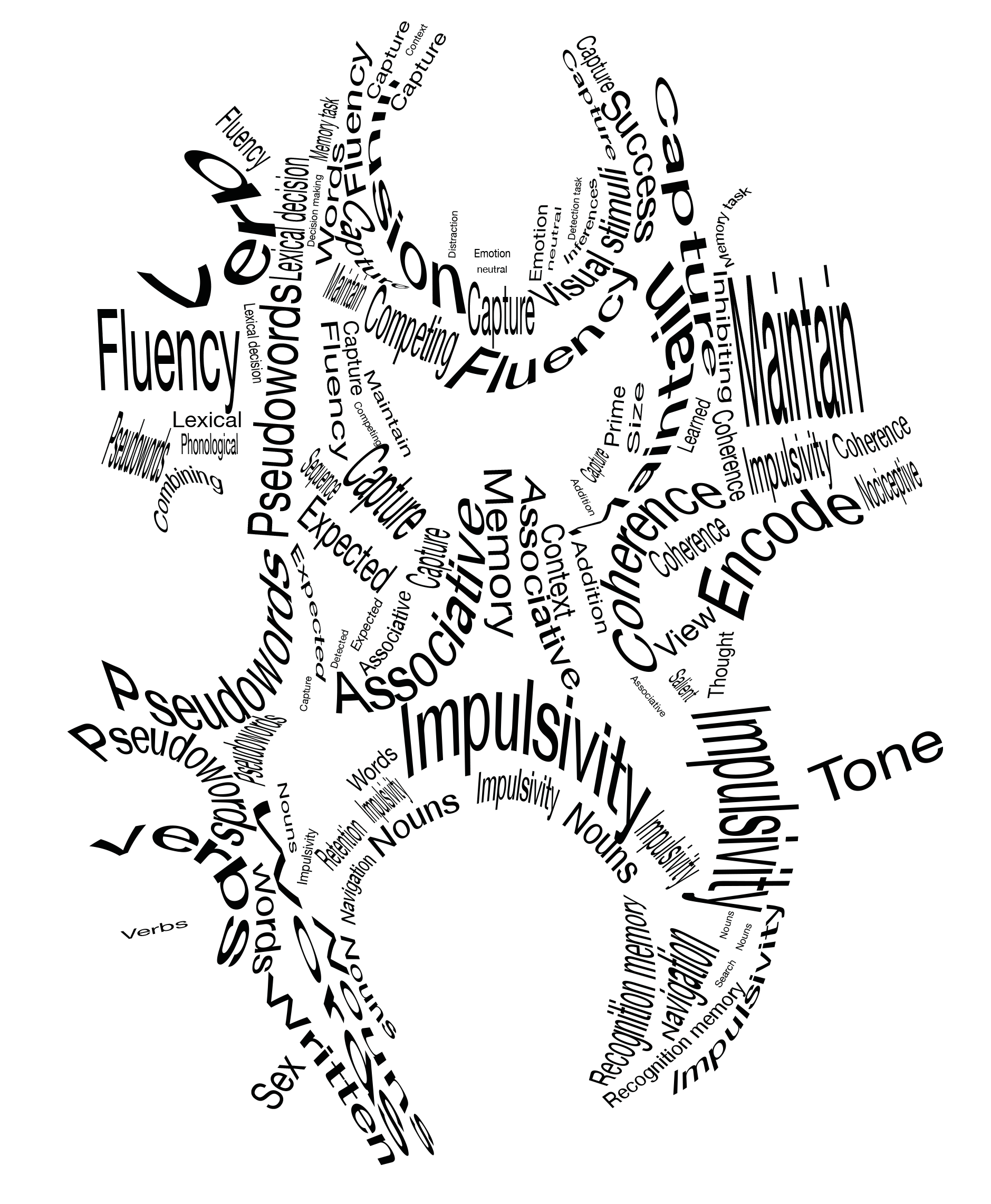

Supplementary figure 52: atlas of white matter function MNI slice z = 20 (neurological convention)

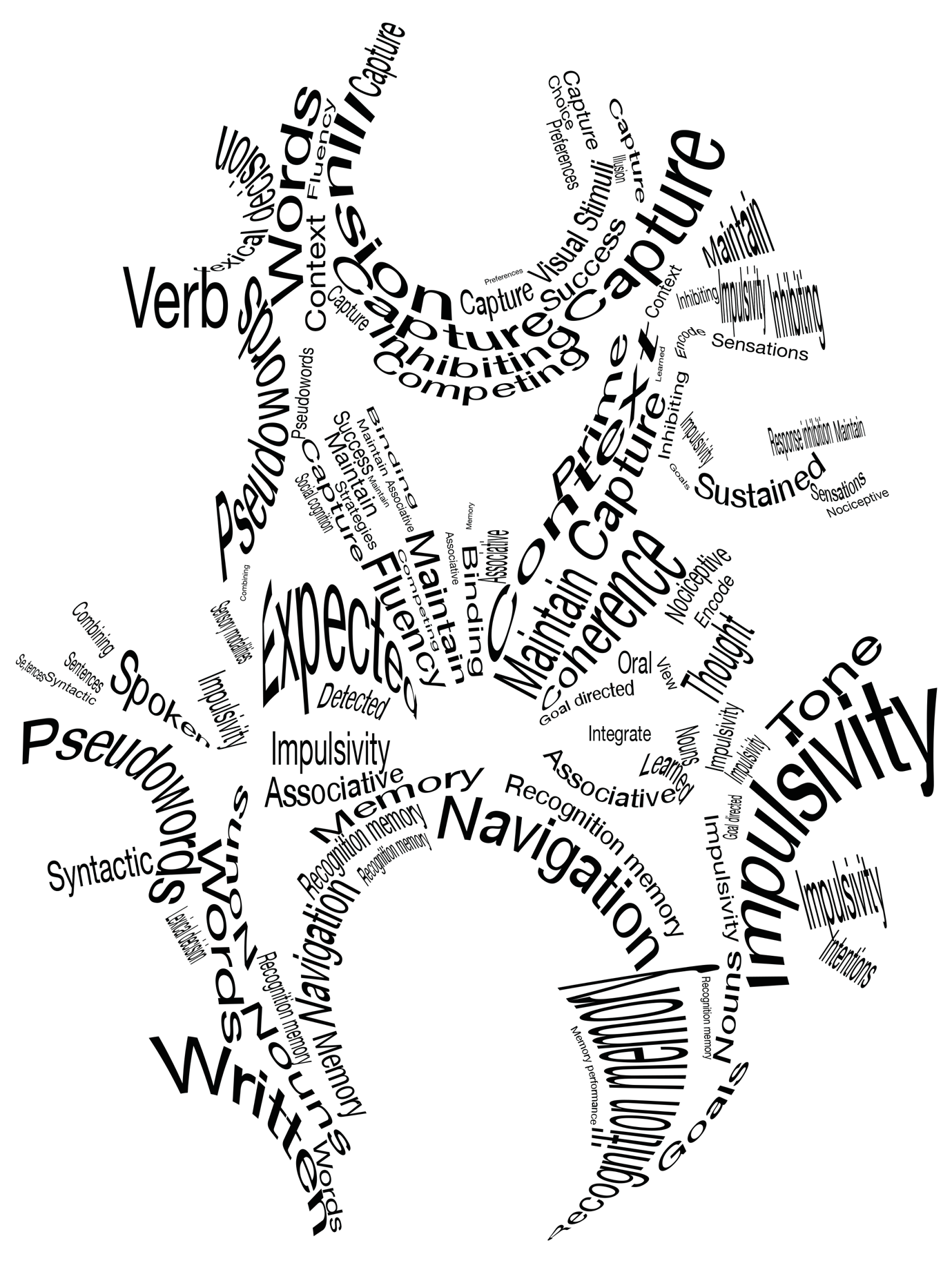

Supplementary figure 53: atlas of white matter function MNI slice z = 10 (neurological convention)

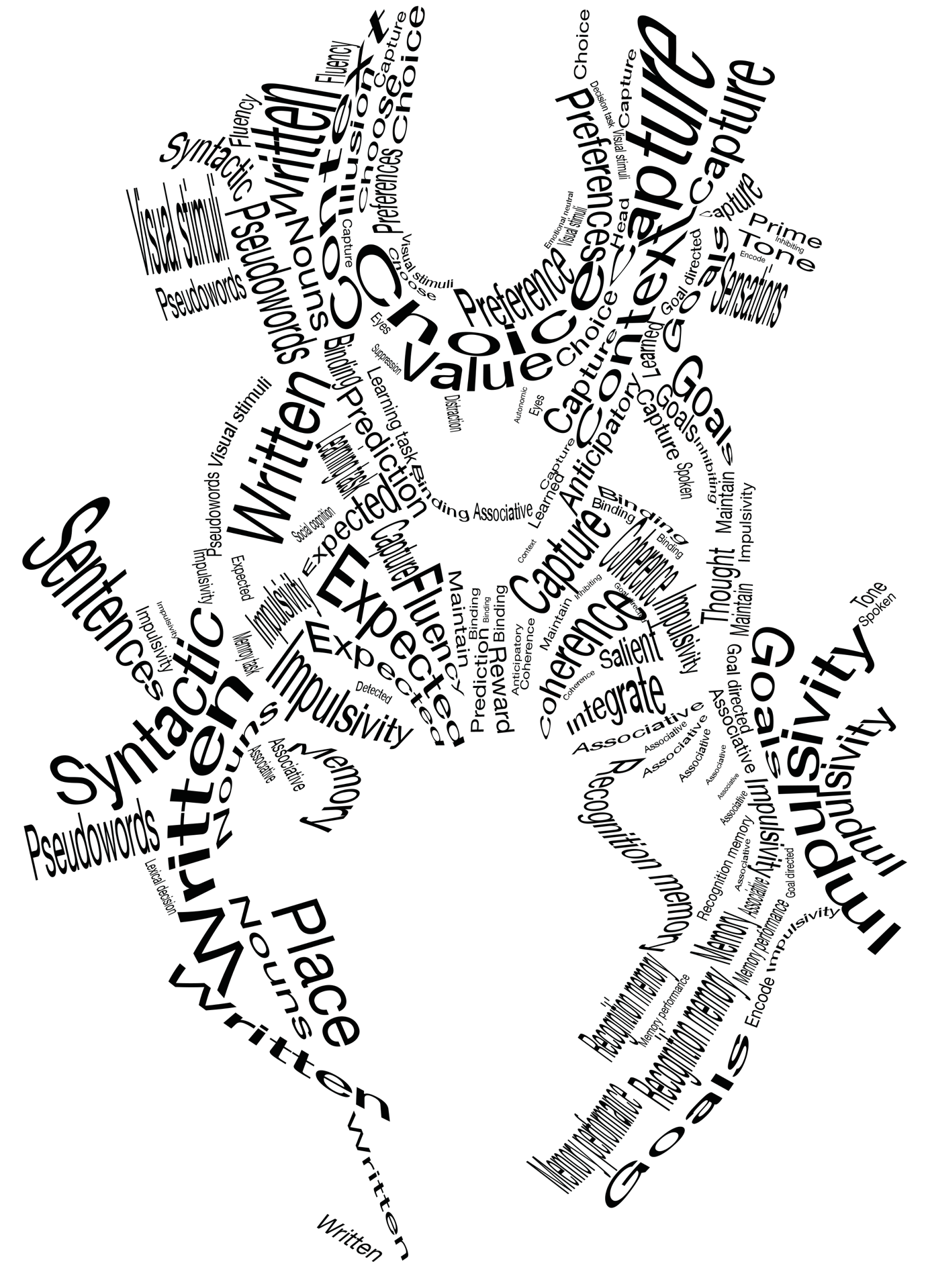

Supplementary figure 54: atlas of white matter function MNI slice z = 0 (neurological convention)

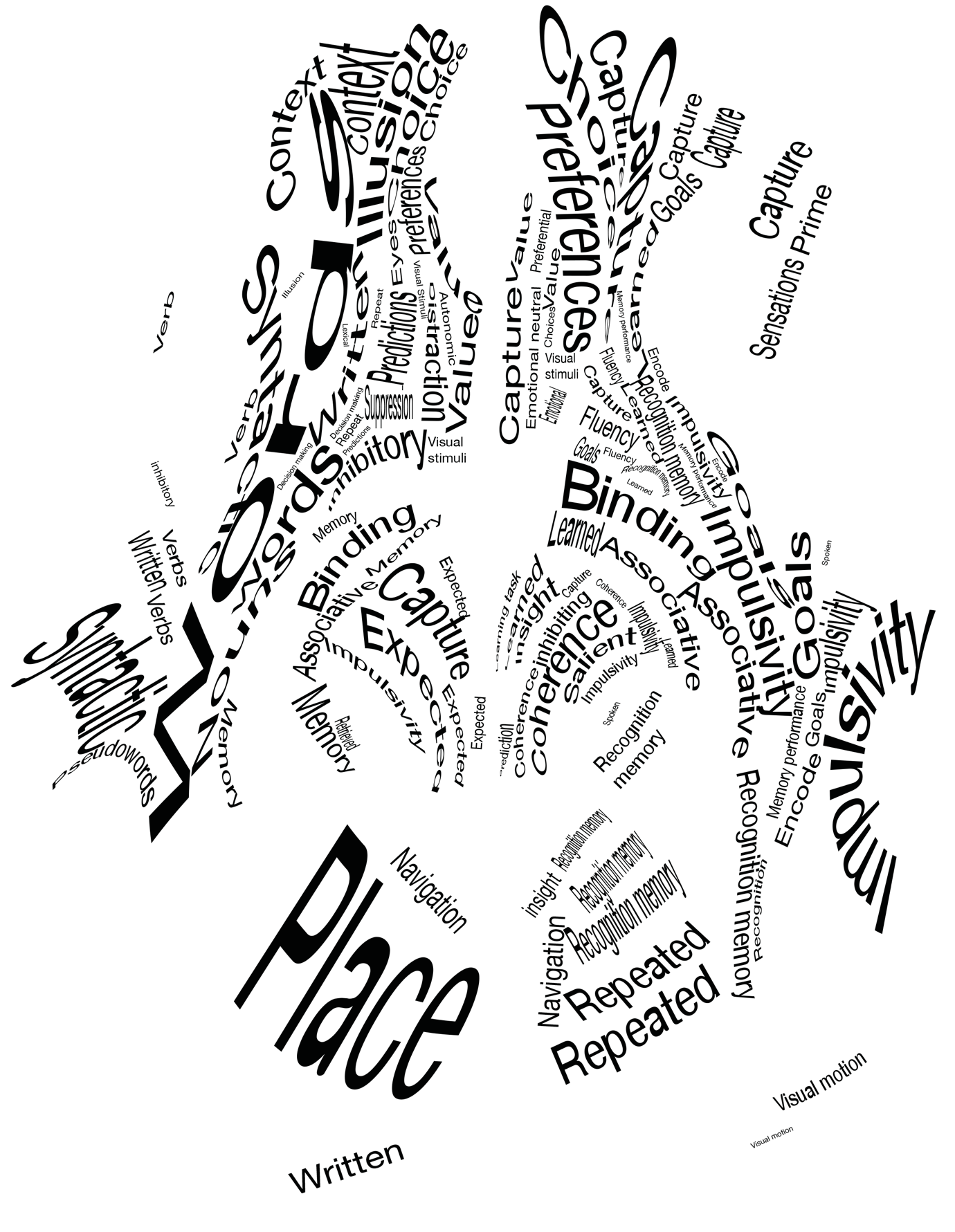

Supplementary figure 54: atlas of white matter function MNI slice z = –10 (neurological convention)

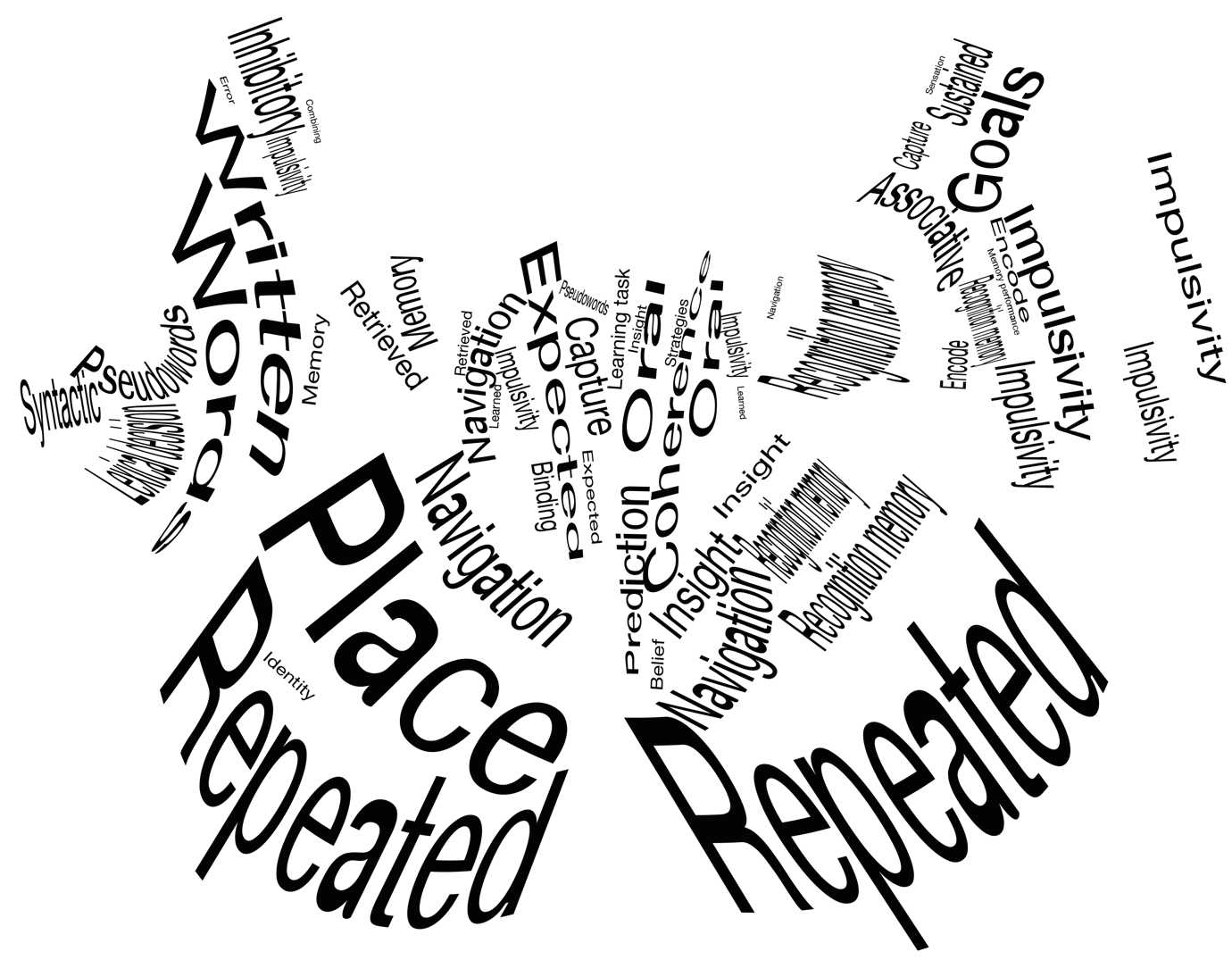

Supplementary figure 55: atlas of white matter function MNI slice z = –20 (neurological convention)

15	O'Keefe, J. & Nadel, L. *The hippocampus as a cognitive map*. (Clarendon Press ;

Oxford University Press, 1978).

16	Mao, D., Kandler, S., McNaughton, B. L. & Bonin, V. Sparse orthogonal population representation of spatial context in the retrosplenial cortex. *Nat Commun* **8**, 243, doi:10.1038/s41467-017-00180-9 (2017).

17	Alexander, A. S. & Nitz, D. A. Spatially Periodic Activation Patterns of Retrosplenial Cortex Encode Route Sub-spaces and Distance Traveled. *Curr Biol* **27**, 1551-1560 e1554, doi:10.1016/j.cub.2017.04.036 (2017).

18	Bussey, T. J. & Saksida, L. M. Memory, perception, and the ventral visual-perirhinal-hippocampal stream: thinking outside of the boxes. *Hippocampus* **17**, 898-908, doi:10.1002/hipo.20320 (2007).

19	Pickens, C. L. *et al.* Different roles for orbitofrontal cortex and basolateral amygdala in a reinforcer devaluation task. *J Neurosci* **23**, 11078-11084 (2003).

20	Zeeb, F. D. & Winstanley, C. A. Functional disconnection of the orbitofrontal cortex and basolateral amygdala impairs acquisition of a rat gambling task and disrupts animals' ability to alter decision-making behavior after reinforcer devaluation. *J Neurosci* **33**, 6434-6443, doi:10.1523/JNEUROSCI.3971-12.2013 (2013).

21	Szpunar, K. K., Watson, J. M. & McDermott, K. B. Neural substrates of envisioning the future. *Proc Natl Acad Sci U S A* **104**, 642-647, doi:10.1073/pnas.0610082104 (2007).

22	Catani, M. & Thiebaut de Schotten, M. *Atlas of Human Brain Connections*. (Oxford University Press, 2012).

23	Yousry, T. A. *et al.* Localization of the motor hand area to a knob on the precentral gyrus. A new landmark. *Brain* **120 ( Pt 1)**, 141-157 (1997).

24	Catani, M. A little man of some importance. *Brain* **140**, 3055-3061, doi:10.1093/brain/awx270 (2017).

25	Akelaitis, A. J. Studies of the corpus callosum: IV. Diagonistic dyspraxia in epileptics following partial and complete section of the corpus callosum. *Am J Psychiat*, 594–599 (1945).

26	Bogen, J. E. in *Surgery of the third ventricle*	(ed Apuzzo M.L.J.) 175–194 (Williams and Wilkins, 1987).

27	Goldstein, K. Zur Lehre von der motorisschen Apraxie. *J Psychol Neurol* **XI**, 169–187 (1908).

28	Della Sala, S., Marchetti, C. & Spinnler, H. Right-sided anarchic (alien) hand: a longitudinal study. *Neuropsychologia* **29**, 1113-1127, doi:10.1016/0028-3932(91)90081-i (1991).

29	Fruhmann Berger, M., Pross, R. D., Ilg, U. & Karnath, H. O. Deviation of eyes and head in acute cerebral stroke. *BMC Neurol* **6**, 23, doi:10.1186/1471-2377-6-23 (2006).

30	Gould, H. J., 3rd, Cusick, C. G., Pons, T. P. & Kaas, J. H. The relationship of corpus callosum connections to electrical stimulation maps of motor, supplementary motor, and the frontal eye fields in owl monkeys. *J Comp Neurol* **247**, 297-325, doi:10.1002/cne.902470303 (1986).

31	Dronkers, N. F., Wilkins, D. P., Van Valin, R. D., Jr., Redfern, B. B. & Jaeger, J. J. Lesion analysis of the brain areas involved in language comprehension. *Cognition* **92**, 145-177, doi:10.1016/j.cognition.2003.11.002

S0010027703002300 [pii] (2004).

32	Mesulam, M. M., Thompson, C. K., Weintraub, S. & Rogalski, E. J. The Wernicke conundrum and the anatomy of language comprehension in primary progressive aphasia. *Brain* **138**, 2423-2437, doi:10.1093/brain/awv154 (2015).

33	Ross, E. D. & Geschwind, N. G. in *The Clinical Neurosciences* Vol. 2 (ed Rosenberg R.N.) 777-796 (Churchill Livingstone, 1983).

34	Kleinschmidt, A. & Rusconi, E. Gerstmann meets Geschwind: a crossing (or kissing) variant of a subcortical disconnection syndrome? *Neuroscientist* **17**, 633-644, doi:10.1177/1073858411402093 (2011).

35	Mayer, E. *et al.* A pure case of Gerstmann syndrome with a subangular lesion. *Brain* **122 ( Pt 6)**, 1107-1120, doi:10.1093/brain/122.6.1107 (1999).

36	Gerstmann, J. Syndrome of finger agnosia, disorientation for right and left, agraphia and acalculia. *Arch Neurol Psychiatry* **44**, 398–407 (1940).

37	Gerstmann, J. Fingeragnosie. Eine umschriebene Störung der Orientierung am eigenen Körper. *Wien Klin Wochenschr* **40**, 1010–1012 (1924).

38	Thiebaut de Schotten, M. *et al.* Direct evidence for a parietal-frontal pathway subserving spatial awareness in humans. *Science* **309**, 2226-2228, doi:10.1126/science.1116251 (2005).

39	Mort, D. J. *et al.* The anatomy of visual neglect. *Brain* **126**, 1986-1997, doi:10.1093/brain/awg200 (2003).

40	Gaffan, D. & Hornak, J. Visual neglect in the monkey. Representation and disconnection. *Brain* **120 ( Pt 9)**, 1647-1657 (1997).

41	Meyer, S. *et al.* Voxel-based lesion-symptom mapping of stroke lesions underlying somatosensory deficits. *Neuroimage Clin* **10**, 257-266, doi:10.1016/j.nicl.2015.12.005 (2016).

42	Catani, M. *et al.* Short frontal lobe connections of the human brain. *Cortex* **48**, 273-291, doi:S0010-9452(11)00317-0 [pii]

10.1016/j.cortex.2011.12.001 (2012).

43	Karnath, H. O., Baier, B. & Nagele, T. Awareness of the functioning of one's own limbs mediated by the insular cortex? *J Neurosci* **25**, 7134-7138, doi:10.1523/JNEUROSCI.1590-05.2005 (2005).

44	Leleux, C., Kaiser, G. & Lebrun, Y. in *Neurolinguistic* Vol. 9 (ed R. Hoops and Y. Lebrun) 141–158 (Sweets and Zeitlinger, 1979).

45	Ardila, A. & Rosselli, M. Acalculia and dyscalculia. *Neuropsychol Rev* **12**, 179-231, doi:10.1023/a:1021343508573 (2002).

46	Grana, A., Hofer, R. & Semenza, C. Acalculia from a right hemisphere lesion dealing with "where" in multiplication procedures. *Neuropsychologia* **44**, 2972-2986, doi:10.1016/j.neuropsychologia.2006.06.027 (2006).

47	Benavides-Varela, S. *et al.* Right-hemisphere (spatial?) acalculia and the influence of neglect. *Front Hum Neurosci* **8**, 644, doi:10.3389/fnhum.2014.00644 (2014).

48	Vaina, L. M., Grzywacz, N. M., LeMay, M., Bienfang, D. & Wolpow, E. in *High-level Motion Processing: Computational, Neurobiological, and Psychophysical Perspectives*	(ed Takeo Watanabe) 211-249 (MIT press, 1998).

49	Vaina, L. M., Cowey, A., Jakab, M. & Kikinis, R. Deficits of motion integration and segregation in patients with unilateral extrastriate lesions. *Brain* **128**, 2134-2145, doi:10.1093/brain/awh573 (2005).

50	Andrews, G. *et al.* Relational processing following stroke. *Brain and Cognition* **81**, 44-51, doi:<https://doi.org/10.1016/j.bandc.2012.09.003> (2013).

51	Bendetowicz, D. *et al.* Two critical brain networks for generation and combination of remote associations. *Brain* **141**, 217-233, doi:10.1093/brain/awx294 (2018).

52	di Pellegrino, G., Ciaramelli, E. & Ladavas, E. The regulation of cognitive control following rostral anterior cingulate cortex lesion in humans. *J Cogn Neurosci* **19**, 275-286, doi:10.1162/jocn.2007.19.2.275 (2007).

53	Thompson-Schill, S. L. *et al.* Verb generation in patients with focal frontal lesions: a neuropsychological test of neuroimaging findings. *Proc Natl Acad Sci U S A* **95**, 15855-15860, doi:10.1073/pnas.95.26.15855 (1998).

54	Foulon, C. *et al.* Advanced lesion symptom mapping analyses and implementation as BCBtoolkit. *Gigascience* **7**, 1-17, doi:10.1093/gigascience/giy004 (2018).

55	Swick, D., Ashley, V. & Turken, A. U. Left inferior frontal gyrus is critical for response inhibition. *BMC Neurosci* **9**, 102, doi:10.1186/1471-2202-9-102 (2008).

56	Nachev, P., Wydell, H., O'Neill, K., Husain, M. & Kennard, C. The role of the pre-supplementary motor area in the control of action. *Neuroimage* **36 Suppl 2**, T155-163, doi:10.1016/j.neuroimage.2007.03.034 (2007).

57	Mandonnet, E. *et al.* A network-level approach of cognitive flexibility impairment after surgery of a right temporo-parietal glioma. *Neurochirurgie* **63**, 308-313, doi:10.1016/j.neuchi.2017.03.003 (2017).

58	Godefroy, O., Lhullier, C. & Rousseaux, M. Non-spatial attention disorders in patients with frontal or posterior brain damage. *Brain* **119 ( Pt 1)**, 191-202, doi:10.1093/brain/119.1.191 (1996).

59	Baldan Ramsey, L. C., Xu, M., Wood, N. & Pittenger, C. Lesions of the dorsomedial striatum disrupt prepulse inhibition. *Neuroscience* **180**, 222-228, doi:10.1016/j.neuroscience.2011.01.041 (2011).

60	Brownell, H. H., Potter, H. H., Bihrle, A. M. & Gardner, H. Inference deficits in right brain-damaged patients. *Brain Lang* **27**, 310-321, doi:10.1016/0093-934x(86)90022-2 (1986).

61	Wapner, W., Hamby, S. & Gardner, H. The role of the right hemisphere in the apprehension of complex linguistic materials. *Brain Lang* **14**, 15-33, doi:10.1016/0093-934x(81)90061-4 (1981).

62	Moya, K. L., Benowitz, L. I., Levine, D. N. & Finklestein, S. Covariant defects in visuospatial abilities and recall of verbal narrative after right hemisphere stroke. *Cortex* **22**, 381-397, doi:10.1016/s0010-9452(86)80003-x (1986).

63	Blake, M. L. Inferencing processes after right hemisphere brain damage: maintenance of inferences. *J Speech Lang Hear Res* **52**, 359-372, doi:10.1044/1092-4388(2009/07-0012) (2009).

64	Silagi, M. L., Radanovic, M., Conforto, A. B., Mendonca, L. I. Z. & Mansur, L. L. Inference comprehension in text reading: Performance of individuals with right- versus left-hemisphere lesions and the influence of cognitive functions. *PLoS One* **13**, e0197195, doi:10.1371/journal.pone.0197195 (2018).

65	Urbanski, M. *et al.* Reasoning by analogy requires the left frontal pole: lesion-deficit mapping and clinical implications. *Brain* **139**, 1783-1799, doi:10.1093/brain/aww072 (2016).

66	Cavanna, A. E. & Trimble, M. R. The precuneus: a review of its functional anatomy and behavioural correlates. *Brain* **129**, 564-583, doi:10.1093/brain/awl004 (2006).

67	Herbet, G. *et al.* Disrupting posterior cingulate connectivity disconnects consciousness from the external environment. *Neuropsychologia* **56**, 239-244, doi:10.1016/j.neuropsychologia.2014.01.020 (2014).

68	Jaynes, J. *The origin of consciousness in the breakdown of the bicameral mind*. (Houghton Mifflin, 1990).

69	Turine, G., Gille, M., Druart, C., Rommel, D. & Rutgers, M. P. Bilateral anterior fornix infarction: the "amnestic syndrome of the subcallosal artery". *Acta Neurol Belg* **116**, 371-373, doi:10.1007/s13760-015-0553-6 (2016).

70	Renou, P., Ducreux, D., Batouche, F. & Denier, C. Pure and acute Korsakoff syndrome due to a bilateral anterior fornix infarction: a diffusion tensor tractography study. *Arch Neurol* **65**, 1252-1253, doi:10.1001/archneur.65.9.1252 (2008).

71	Park, S. A., Hahn, J. H., Kim, J. I., Na, D. L. & Huh, K. Memory deficits after bilateral anterior fornix infarction. *Neurology* **54**, 1379-1382, doi:10.1212/wnl.54.6.1379 (2000).

72	Moudgil, S. S., Azzouz, M., Al-Azzaz, A., Haut, M. & Gutmann, L. Amnesia due to fornix infarction. *Stroke* **31**, 1418-1419, doi:10.1161/01.str.31.6.1418 (2000).

73	Kawada, T. Isolated bilateral fornix stroke and acute amnestic syndrome. *Eur J Neurol* **26**, e63, doi:10.1111/ene.13849 (2019).

74	Salvalaggio, A., Cagnin, A., Nardetto, L., Manara, R. & Briani, C. Acute amnestic syndrome in isolated bilateral fornix stroke. *Eur J Neurol* **25**, 787-789, doi:10.1111/ene.13592 (2018).

75	Aggleton, J. P. EPS Mid-Career Award 2006. Understanding anterograde amnesia: disconnections and hidden lesions. *Q J Exp Psychol (Hove)* **61**, 1441-1471, doi:10.1080/17470210802215335 (2008).

76	Husain, M. & Nachev, P. Space and the parietal cortex. *Trends Cogn Sci* **11**, 30-36, doi:10.1016/j.tics.2006.10.011 (2007).

77	Hayashi, M. J., van der Zwaag, W., Bueti, D. & Kanai, R. Representations of time in human frontoparietal cortex. *Commun Biol* **1**, 233, doi:10.1038/s42003-018-0243-z (2018).

78	Verdon, V., Schwartz, S., Lovblad, K. O., Hauert, C. A. & Vuilleumier, P. Neuroanatomy of hemispatial neglect and its functional components: a study using voxel-based lesion-symptom mapping. *Brain* **133**, 880-894, doi:10.1093/brain/awp305 (2010).

79	Bartolomeo, P., Thiebaut de Schotten, M. & Doricchi, F. Left unilateral neglect as a disconnection syndrome. *Cereb Cortex* **17**, 2479-2490, doi:10.1093/cercor/bhl181 (2007).

80	Ghika, J., Bogousslavsky, J., Uske, A. & Regli, F. Parietal kinetic ataxia without proprioceptive deficit. *J Neurol Neurosurg Psychiatry* **59**, 531-533, doi:10.1136/jnnp.59.5.531 (1995).

81	Preilowski, B. F. Possible contribution of the anterior forebrain commissures to bilateral motor coordination. *Neuropsychologia* **10**, 267-277, doi:10.1016/0028-3932(72)90018-8 (1972).

82	Eliassen, J. C., Baynes, K. & Gazzaniga, M. S. Anterior and posterior callosal contributions to simultaneous bimanual movements of the hands and fingers. *Brain* **123 Pt 12**, 2501-2511, doi:10.1093/brain/123.12.2501 (2000).

83	Behrman, M. & Kimchi, R. What does visual agnosia tell us about perceptual organization and its relationship to object perception? *Journal of Experimental Psychology: Human Perception and Performance* **42**, 19-42 (2003).

84	Riddoch, M. J. & Humphreys, G. W. A case of integrative visual agnosia. *Brain* **110 ( Pt 6)**, 1431-1462 (1987).

85	Riddoch, M. J., Johnston, R. A., Bracewell, R. M., Boutsen, L. & Humphreys, G. W. Are faces special? A case of pure prosopagnosia. *Cogn Neuropsychol* **25**, 3-26, doi:791466789 [pii]

10.1080/02643290801920113 (2008).

86	Ferreira, C. T., Ceccaldi, M., Giusiano, B. & Poncet, M. Separate visual pathways for perception of actions and objects: evidence from a case of apperceptive agnosia. *Journal of neurology, neurosurgery, and psychiatry* **65**, 382-385 (1998).

87	Warrington, E. K. & James, M. Visual apperceptive agnosia: a clinico-anatomical study of three cases. *Cortex; a journal devoted to the study of the nervous system and behavior* **24**, 13-32 (1988).

88	Jankowiak, J. & Albert, M. L. *Lesion localization in visual agnosia. In: Kertesz A, editor. Localization and neuroimaging in neuropsychology*. 429-471 (Academic Press, 1994).

89	Harlow, J. Passage of an iron bar through the head. *Boston Med. Surg. J.* **39**, 389–393 (1848).

90	Thiebaut de Schotten, M. *et al.* From Phineas Gage and Monsieur Leborgne to H.M.: Revisiting Disconnection Syndromes. *Cereb Cortex* **25**, 4812-4827, doi:10.1093/cercor/bhv173 (2015).

91	Thiebaut de Schotten, M., Dell'acqua, F., Valabregue, R. & Catani, M. Monkey to human comparative anatomy of the frontal lobe association tracts. *Cortex* **48**, 82-96, doi:S0010-9452(11)00275-9 [pii]

10.1016/j.cortex.2011.10.001 (2012).

92	Dronkers, N. F., Plaisant, O., Iba-Zizen, M. T. & Cabanis, E. A. Paul Broca's historic cases: high resolution MR imaging of the brains of Leborgne and Lelong. *Brain* **130**, 1432-1441 (2007).

93	Turken, A. U. & Dronkers, N. F. The neural architecture of the language comprehension network: converging evidence from lesion and connectivity analyses. *Front Syst Neurosci* **5**, 1, doi:10.3389/fnsys.2011.00001 (2011).

94	Snaphaan, L., Rijpkema, M., van Uden, I., Fernandez, G. & de Leeuw, F. E. Reduced medial temporal lobe functionality in stroke patients: a functional magnetic resonance imaging study. *Brain* **132**, 1882-1888, doi:10.1093/brain/awp133 (2009).

95	Berg, E. A. A simple objective technique for measuring flexibility in thinking. *J. Gen. Psychol.* **39**, 15-22 (1948).

96	Glascher, J., Adolphs, R. & Tranel, D. Model-based lesion mapping of cognitive control using the Wisconsin Card Sorting Test. *Nat Commun* **10**, 20, doi:10.1038/s41467-018-07912-5 (2019).

97	Gabrieli, J. D. E., Fleischman, D. A., Keane, M. M., Reminger, S. L. & Morrell, F. Double Dissociation between Memory Systems Underlying Explicit and Implicit Memory in

the Human Brain. *Psychological Science* **6**, 76-82 (1995).

98	Bird, C. M. *et al.* Visual neglect after right posterior cerebral artery infarction. *J Neurol Neurosurg Psychiatry* **77**, 1008-1012, doi:10.1136/jnnp.2006.094417 (2006).

99	Maffei, C., Sarubbo, S. & Jovicich, J. A Missing Connection: A Review of the Macrostructural Anatomy and Tractography of the Acoustic Radiation. *Front Neuroanat* **13**, 27, doi:10.3389/fnana.2019.00027 (2019).

100	Sihvonen, A. J. *et al.* Neural Basis of Acquired Amusia and Its Recovery after Stroke. *J Neurosci* **36**, 8872-8881, doi:10.1523/JNEUROSCI.0709-16.2016 (2016).

101	Sihvonen, A. J., Ripolles, P., Rodriguez-Fornells, A., Soinila, S. & Sarkamo, T. Revisiting the Neural Basis of Acquired Amusia: Lesion Patterns and Structural Changes Underlying Amusia Recovery. *Front Neurosci* **11**, 426, doi:10.3389/fnins.2017.00426 (2017).

102	Jafari, Z., Esmaili, M., Delbari, A., Mehrpour, M. & Mohajerani, M. H. Post-stroke acquired amusia: A comparison between right- and left-brain hemispheric damages. *NeuroRehabilitation* **40**, 233-241, doi:10.3233/NRE-161408 (2017).

103	Karnath, H. O., Ferber, S. & Himmelbach, M. Spatial awareness is a function of the temporal not the posterior parietal lobe. *Nature* **411**, 950-953, doi:10.1038/35082075 (2001).

104	Corbetta, M., Kincade, M. J., Lewis, C., Snyder, A. Z. & Sapir, A. Neural basis and recovery of spatial attention deficits in spatial neglect. *Nature Neuroscience* **8**, 1603-1610 (2005).

105	Beis, J. M. *et al.* Right spatial neglect after left hemisphere stroke: qualitative and quantitative study. *Neurology* **63**, 1600-1605 (2004).

106	Hausler, R. & Levine, R. A. Auditory dysfunction in stroke. *Acta Otolaryngol* **120**, 689-703, doi:10.1080/000164800750000207 (2000).

107	Tolomeo, S. *et al.* A causal role for the anterior mid-cingulate cortex in negative affect and cognitive control. *Brain* **139**, 1844-1854, doi:10.1093/brain/aww069 (2016).

108	Ciaramelli, E., Sperotto, R. G., Mattioli, F. & di Pellegrino, G. Damage to the ventromedial prefrontal cortex reduces interpersonal disgust. *Soc Cogn Affect Neurosci* **8**, 171-180, doi:10.1093/scan/nss087 (2013).

109	Ho, C. Y. & Berridge, K. C. Excessive disgust caused by brain lesions or temporary inactivations: mapping hotspots of the nucleus accumbens and ventral pallidum. *Eur J Neurosci* **40**, 3556-3572, doi:10.1111/ejn.12720 (2014).

110	Tzourio-Mazoyer, N. *et al.* Between-hand difference in ipsilateral deactivation is associated with hand lateralization: fMRI mapping of 284 volunteers balanced for handedness. *Front Hum Neurosci* **9**, 5, doi:10.3389/fnhum.2015.00005 (2015).
