## Supplementary figures and images for "Brain disconnections link structural connectivity with function and behaviour"

### supplementary figure 1

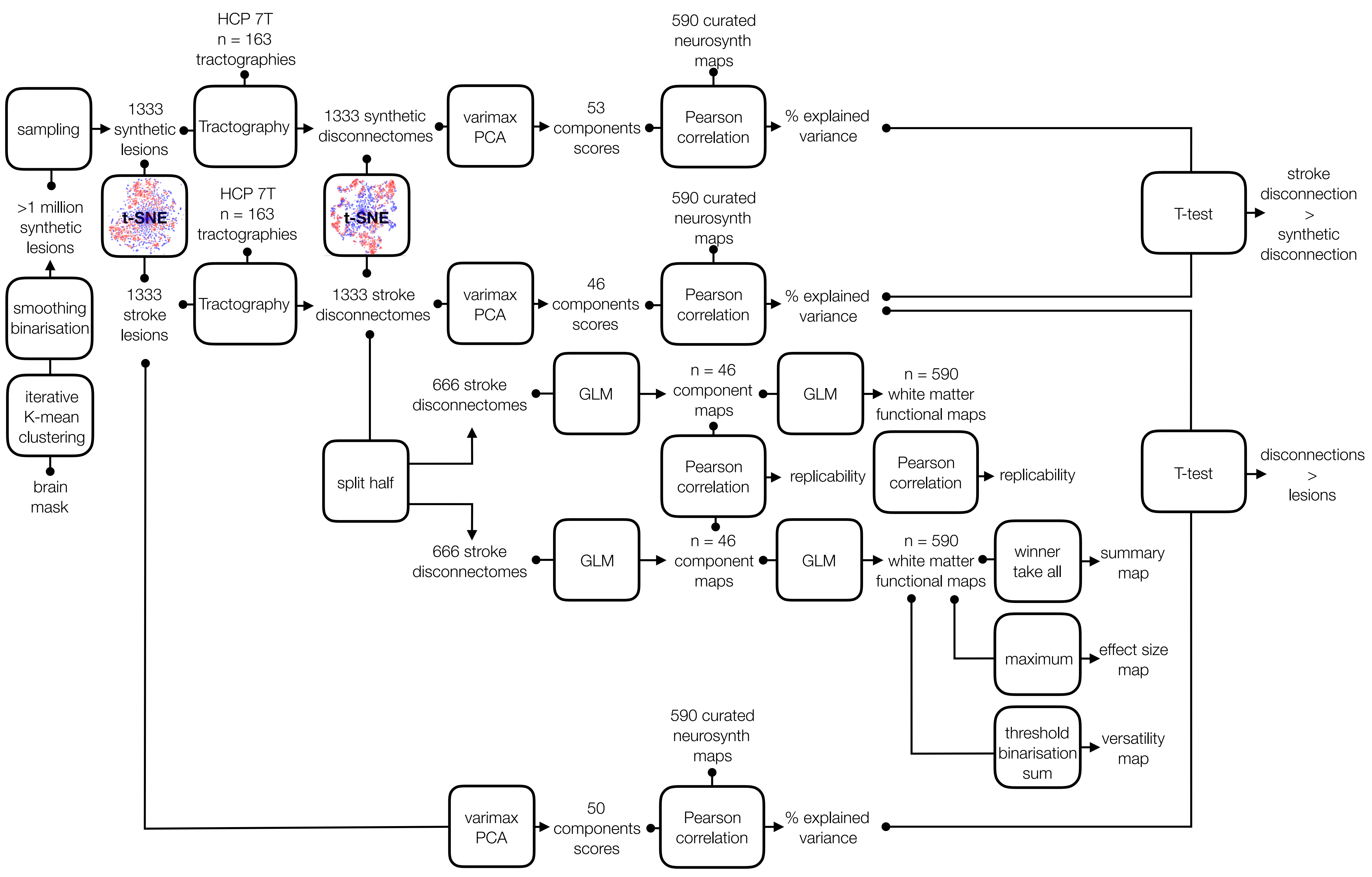
